# Expanding luciferase reporter systems for cell-free protein expression

**DOI:** 10.1101/2022.05.10.491427

**Authors:** Wakana Sato, Melanie Rasmussen, Christopher Deich, Aaron E. Engelhart, Katarzyna P. Adamala

## Abstract

Luciferases are often used as a sensitive, versatile reporter in cell-free transcription-translation (TXTL) systems, for research and practical applications such as engineering genetic parts, validating genetic circuits, and biosensor outputs. Currently, only two luciferases (Firefly and Renilla) are commonly used without substrate cross-talk. Here we demonstrate expansion of the cell-free luciferase reporter system, with two orthogonal luciferase reporters: *N. nambi* luciferase (Luz) and LuxAB. These luciferases do not have cross-reactivity with the Firefly and Renilla substrates. We also demonstrate a substrate regeneration pathway for one of the new luciferases, enabling long-term time courses of protein expression monitoring in the cell-free system. Furthermore, we reduced the number of genes required in TXTL expression, by engineering a cell extract containing part of the luciferase enzymes. Our findings lead to an expanded platform with multiple orthogonal luminescence translation readouts for *in vitro* protein expression.

## Introduction

The cell-free transcription-translation (TXTL) is a widely used *in vitro* protein expression system for synthetic biology^1–3^. *E. coli*-based TXTL has been expanding its usage with intensive engineering efforts^4– 8^. By coupling with reporter genes, TXTL can be used for many applications, such as viral detection^9^, metabolic modeling^10^, toxin detection^11^, biosensors^12^, and genetic circuit validation^13^. In TXTL, the most common reporter genes are luciferases and fluorescent proteins. While luciferases have a higher signal-to-noise ratio than fluorescence proteins^11^, they cannot be used to measure long-term kinetics, due to the nature of the flash reaction substrate-dependency. Thus, fluorescent proteins are preferred for measuring gene expression dynamics. Furthermore, both fluorescent proteins and luciferases are limited for their multiplexing capacity. The fluorescent proteins are limited to about four to five colors for a simultaneous measurement due to their broad emission spectra^14^. As for luciferases, although emission filters allow multiple measurements up to around six^15,16^, the number of available substrates without cross-reactivity is low. The most commonly used substrates are D-luciferin (for Firefly luciferase) and coelenterazine (for Renilla luciferase). Vargulin was recently reported for an additional no-cross-reactive substrate with *Cypridina* luciferase^17^.

Here we address two needs of the luciferase reporter systems in TXTL: expanding multiplexing capabilities, and enabling long-term kinetics measurements. We also demonstrate a TXTL extract preparation that enables the use of luciferase with minimal burden on TXTL resources.

We explored luciferase variants without substrate cross-reactivity, to construct luciferase pathways independent of substrate supplementation, and to optimize their reactions for TXTL. We focused on two luciferase systems: fungi- and bacterial-luciferases. Neither of those luciferases was previously used in TXTL, and both are capable of substrate regeneration^18–23^.

In the fungi luciferase system, hispidin is converted to 3-hydroxyhispidin by *Neonothopanus nambi* (*N. nambi*) hispidin-3-hidroxylase (H3H), and then *N. nambi* luciferase (Luz) yields light by reacting with 3-hydroxyhispidin. Caffeylpyruvate hydrolase (CPH), hispidin synthase (Hisps), and 4’-phosphopantetheinyl transferase (NPGA) are also required for the substrate regeneration^18–20^ (**Fig. 1A, Fig. S10**). The bacterial candidate, LuxABCDE, is a well-characterized and widely used luciferase^21–23^. LuxAB is a luciferase complex that yields light by converting reduced flavin (FMNH_2_) and long-chain aldehydes into oxidized flavin mononucleotide (FMN) and the corresponding long-chain acids. LuxCDE reduces the long-chain acids back to the long-chain aldehydes to regenerate their substrate (**Fig. 1C**).

**Figure 1.**
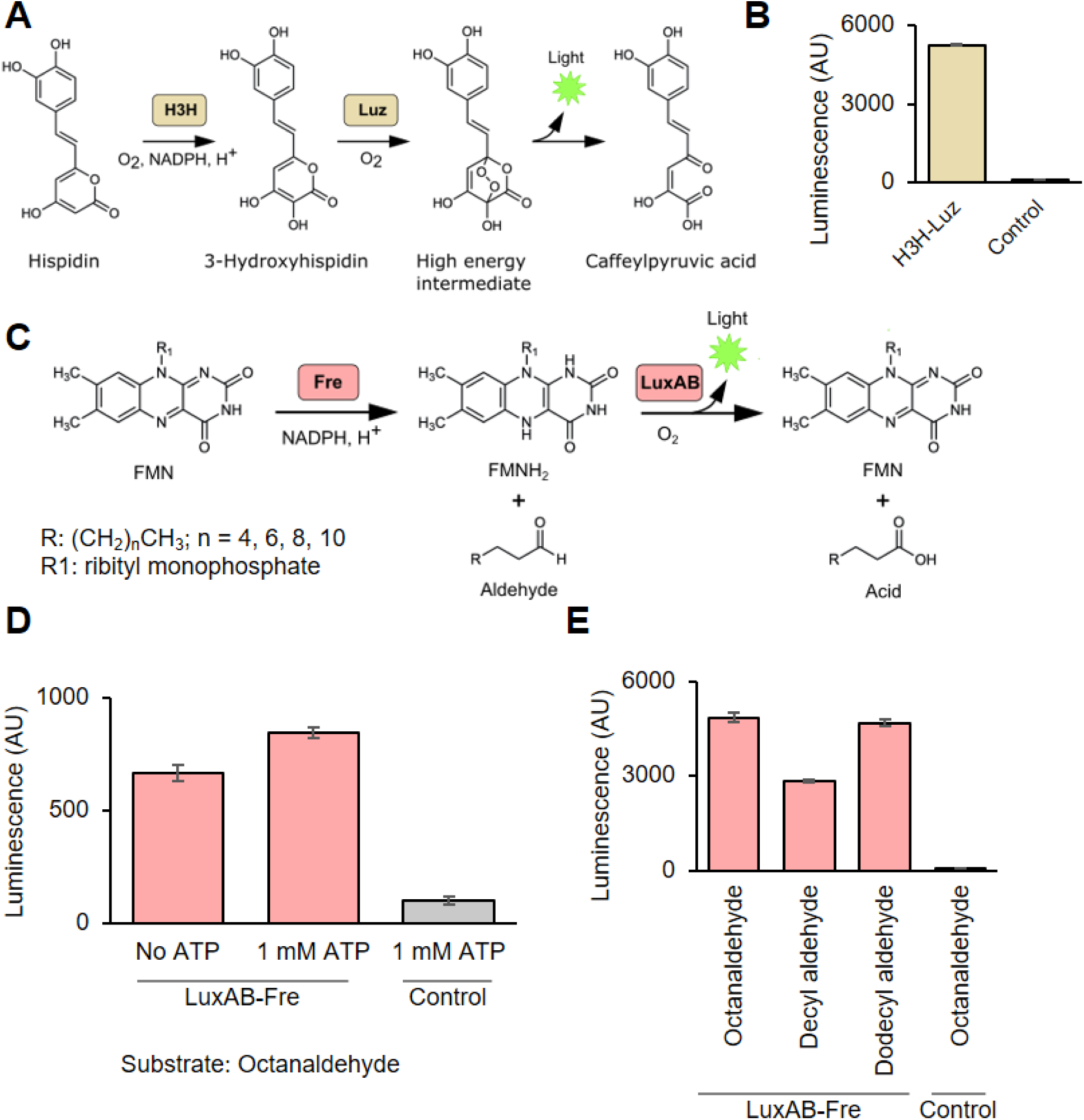
Characterization of H3H-Lux and LuxAB-Fre luciferase systems in TXTL. (A) Schematic of H3H-Luz luciferase reaction. Hispidin is converted to 3-hydroxyhispidin by hispidin-3-hydroxylase (H3H) and 3-hydroxyhispidin is oxidized and converted into a high energy intermediate by the luciferase (Luz). This intermediate decays into caffeylpyruvic acid with light emission. (B) The H3H-Luz luminescence measurement. H3H and Luz were expressed in TXTL. The luminescence was measured right after adding NADPH and hisipidin into the TXTL. (C) Schematic of LuxAB-Fre luciferase reaction. Oxidized flavin mononucleotide (FMN) is reduced into reduced flavin mononucleotide (FMNH_2_) by NAD(P)H-flavin reductase (Fre). The luciferase (LuxAB) converts FMNH_2_ and long-chain aldehydes into FMN and the corresponding long-chain acids with light emission. (E) ATP supplementation increased the light emission of LuxAB-Fre. Octanaldehyde was added as the substrate. (D) The LuxAB-Fre luminescence measurement with different long-chain fatty aldehydes. LuxA, LuxB and Fre were expressed in TXTL. The luminescence was measured right after adding FMN, NADPH, ATP, and substrates (octanaldehyde, decyl aldehyde, and dodecyl aldehyde.) NADPH, nicotinamide adenine dinucleotide phosphate; ATP, adenosine triphosphate; Control, reaction without enzyme expression. The graphs show means with error bars that signify SEM (n = 3).

## Results

Here we report a successful demonstration of using those two luciferase-substrate systems in bacterial TXTL. The fungi luciferase (H3H-Luz) consists of Luz and H3H and generates light from hispidin, a commercially available chemical. The bacterial luciferase (LuxAB-Fre) consists of LuxAB and NAD(P)H-flavin reductase (Fre).

### New luciferase systems for TXTL

We first tested the H3H-Luz and LuxAB-Fre luciferase activities in TXTL. To our knowledge, neither of the reaction has been reported in TXTL. H3H-Luz is a fungi-originated luciferase system that uses hispidin as substrate (**Fig. 1A**)^18–20^. LuxAB is a bacterial luciferase that uses long-chain fatty aldehyde and reduced flavin mononucleotide (FMNH_2_) to generate light^21^. We combined LuxAB with NAD(P)H-flavin reductase (Fre) to reduce FMN back to FMNH_2_ **(Fig. 1C)**^22,24^. To test their activities in TXTL, we individually cloned H3H, Luz, LuxA, LuxB, and Fre genes into a vector plasmid designed for T7 RNA polymerase-coupled TXTL expression^25^. We confirmed that both Luz-H3H and LuxAB-Fre systems generate luminescence **(Fig. 1B** and **D)**. In addition to the plasmids, we added hispidin and NADPH into the H3H-Luz luciferase reaction; octanaldehyde, FMN, and NADPH into the LuxAB-Fre luciferase reaction. Based on the mechanism known for LuxAB, ATP is not the essential compound^21,24^; however, we found ATP enhances luminescence **(Fig. 1D)**. Thus, we also supplemented ATP for the later reactions of LuxAB-Fre system in this report. Additionally, we tested three different long-chain fatty aldehydes with LuxAB-Fre system and confirmed that all generate luminescence **(Fig. 1E)**. Since octanaldehyde showed a strong luminescence and high solubility in the reaction, we chose octanaldehyde as the standard substrate in this report.

### Substrate specificities among different luciferase systems

Next, we examined the substrate specificities of H3H-Luz and LuxAB-Fre by comparing the widely used luciferase-substrate pairs: Firefly luciferase (FLuc)-D-luciferin, Renilla luciferase (RLuc)-coelenterazine h, and Nano luciferase (NanoLuc)-furimazine (**Fig. 2A**). First, we tested those substrates individually. Except for NanoLuc, all the luciferases showed significantly stronger luminescence with the substrate supposed to react than others (**Fig. 2B**). Then, we prepared all the four substrates mixture (All mixture: D-luciferin, coelenterazine h, hispidin, and octanaldehyde). We tested each luciferase to see differences in the luminescence with “All” or “All minus one,” which is without a suitable substrate. We omitted the NanoLuc-furimazine pair from this experiment because NanoLuc reacted with both coelenterazine h and furimazine (**Fig. 2B**). FLuc, RLuc, H3H-Luz, and LuxAB-Fre showed expected luminescence patterns; only the “All mixture” luminesced and the “All minus one” did not (**Fig. 2C-F**). Note that FLuc showed slight luminescence with the “All minus one” because of the reactivity with coelenterazine h (**Fig. S2**). Because the signal-ratio intensity difference was significant, we considered it the success of differentiation.

**Figure 2.**
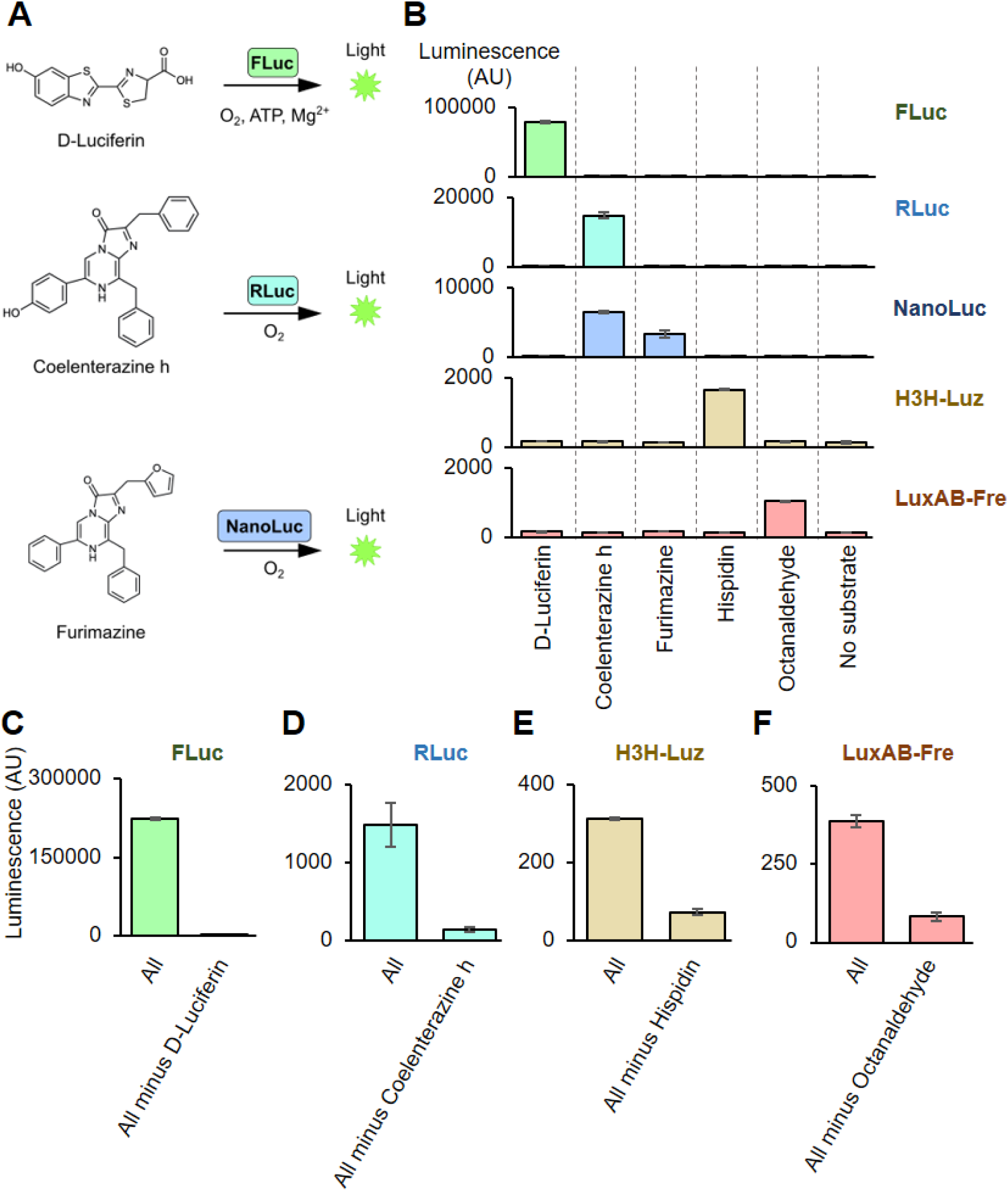
Characterization of substrate specificities. (A) Schematic image of firefly luciferase (FLuc), renilla luciferase (RLuc), and NanoLuc luciferase (NanoLuc) reactions. FLuc oxidizes D-luciferin with ATP and Mg^+^ to produce light. RLuc and NanoLuc oxidize coelenterazine h and furimazine, respectively, with ATP to produce light. (B) Luminescence measurement for substrate specificity assay for 5 luciferases. The luciferases (FLuc, RLuc, NanoLuc, H3H-Luz, and LuxAB-Fre) were expressed in TXTL. Then, the individual substrates (D-luciferin, coelenterazine h, furimazine, hispidin, and octanaldehyde) with corresponding co-factors were added to the reaction and measured its light emission without emission filters. Substrate concentrations were 10 μM, except 1 mM for octanaldehyde. (C-F) The substrate multiplexing assay. The substrate mixtures were prepared as “All” (D-luciferin, Coelenterazine h, hispidin, octanaldehyde, Mg^+^, ATP, NADPH, FMN) or “All minus one” that contains all except one that a substrate is supposed to react with a tested luciferase. The assay was performed by mixing substrates with TXTL expressing (C) FLuc, (D) RLuc, (E) H3H-Luz, or (F) LuxAB-Fre, and the luminescence was measured without emission filters. ATP, adenosine triphosphate. The graphs show means with error bars that signify SEM (n = 3).

### Burden reduced TXTL and fusion proteins with H3H-Luz system

H3H-Luz and LuxAB-Fre systems require multiple gene expression, which may add an extra metabolic burden to TXTL compared to other single gene luciferases (FLuc, RLuc, and NanoLuc). In general, TXTL reactions with fewer genes produce more definite results. Thus, we prepared a cell-free extract containing H3H in advance; we only needed to express Luz in TXTL for luminescence. We used *E. coli* carrying H3H plasmid in the cell-free extract preparation (**Fig. 3A**). This plasmid encodes the H3H gene under the sigma 70 promoter, *E. coli* constitutive promoter. We confirmed that H3H pre-expressed TXTL luminesced as Luz expressed in the presence of hispidin, while the minus Luz reaction did not (**Fig 3B**). We also tested if Luz can be used as a fusion protein. We prepared two Luz-fusion constructs: His-eGFP-Luz and Luz-eGFP-His (described in N-term to C-term order.) 2x GS-linkers (GGGGS) were inserted between the eGFP and Luz genes. Both N-term and C-term Luz fusion constructs luminesced, although the signal intensities were lower than the Luz without fusion proteins (**Fig. 3C**). eGFP fluorescence was stronger for N-term eGFP fusion when the fusion Luz was expressed in TXTL (**Fig. 3D**). Altogether, we claim that a protein can be fused on either terminal of Luz; however, fusing a protein of interest in the N-term of Luz might work better, based on the eGFP fluorescence measurement. We also tried these burden reduction TXTL and fusion protein experiments with the LuxAB-Fre system; however, we failed. We could not detect any luminescence with the *E. coli* strain carrying all the enzymes except LuxA or LuxB (data not shown). Most fusion constructs did not fluoresce or luminesce (**Fig. S4**).

**Figure 3.**
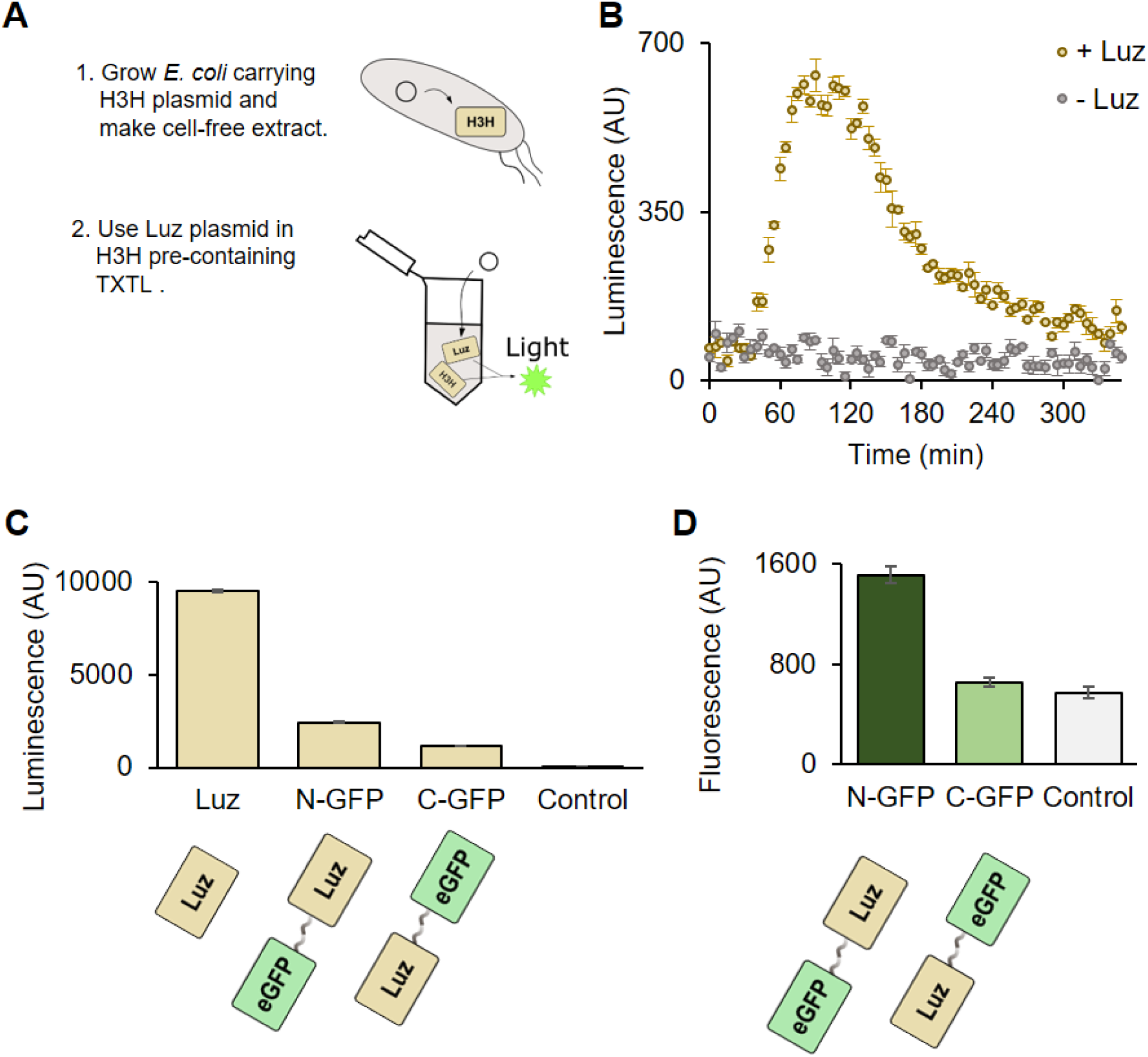
H3H-Luz tested with burden-reduced TXTL and fusion proteins. (A) The schematic of how H3H-carrying TXTL works. The plasmid coding H3H gene under the sigma 70 promoter is transformed into *E. coli* Rosseta 2 strain. The cell-free extract is made with that strain; thus, the extract contains H3H. Once Luz is expressed in the TXTL, Luz produces luminescence by coordinating with H3H. (B) Luminescence measurement in the H3H pre-containing TXTL. Luz plasmids were incubated with hispidin at 30°C. The Luz plasmid containing reaction (yellow dots) generated light during the TXTL reaction, while the reaction without Luz plasmid did not (black squares.) (C) Luminescence measurement of the H3H-Luz system with eGFP fused Luz constructs. H3H and Luz proteins were expressed in TXTL. After the expression, hispidin and NADPH were added, followed by luminescence measurement. (D) Fluorescence measurement of the eGFP fused Luz. Luz proteins were expressed in TXTL and the fluorescence was measured. Luz, Luz luciferase without a fusion protein; N-GFP, N-terminal eGFP fusion with Luz; C-GFP, C-terminal eGFP fusion with Luz; Control, reaction without enzyme expression. The graphs show means with error bars that signify SEM (n = 3).

### Substrate regeneration with LuxABCDE-Fre system

We propose that LuxAB-Fre system be used for continuous luminescence reaction, although the LuxAB-Fre system did not work well for the metabolic burden reduction TXTL and fusion protein experiments. LuxCDE, a protein complex encoded in another part of the LuxABCDE operon, reduces long-chain fatty acids into corresponding long-chain fatty aldehydes^21^. After the luminescence, Fre and LuxCDE can convert the FMN and long-chain fatty acids back to the LuxAB substrates. Thus, LuxABCDE-Fre can self-replenish its substrates consistently **(Fig. 4A)**. First, we tested the luminescence production from long-chain fatty acids. This reaction requires LuxCDE to reduce long-chain fatty acids to long-chain aldehydes. All the long-chain fatty acids we tried (octanoic acid, decanoic acid, dodecanoic acid, and tetradecanoic acid) gave luminescence **(Fig. 4B)**. We set the reaction with octanaldehyde as a positive control to ensure the LuxAB-Fre works **(Fig. 4B)**. Next, we tested whether the LuxABCDE-Fre system can regenerate the substrate to give continuous luminescence. We added the substrate (decanoic acid or octanaldehyde) into the TXTL that had expressed LuxAB-Fre and LuxCDE to incubate at 25°C. The luminescence was measured at 0, 0.5, 1, 6, and 8 hours. Only the reaction containing all the LuxABCDE-Fre enzymes retained the luminescence over 8 hours **(Fig. 4C, Fig. S7)**. We also tried to reconstitute a substrate regenerative fungi luciferase system; however, we failed with our TXTL system (Supplemental information section, “Efforts to engineer the fungi luciferase substrate regeneration pathway.”)

**Figure 4.**
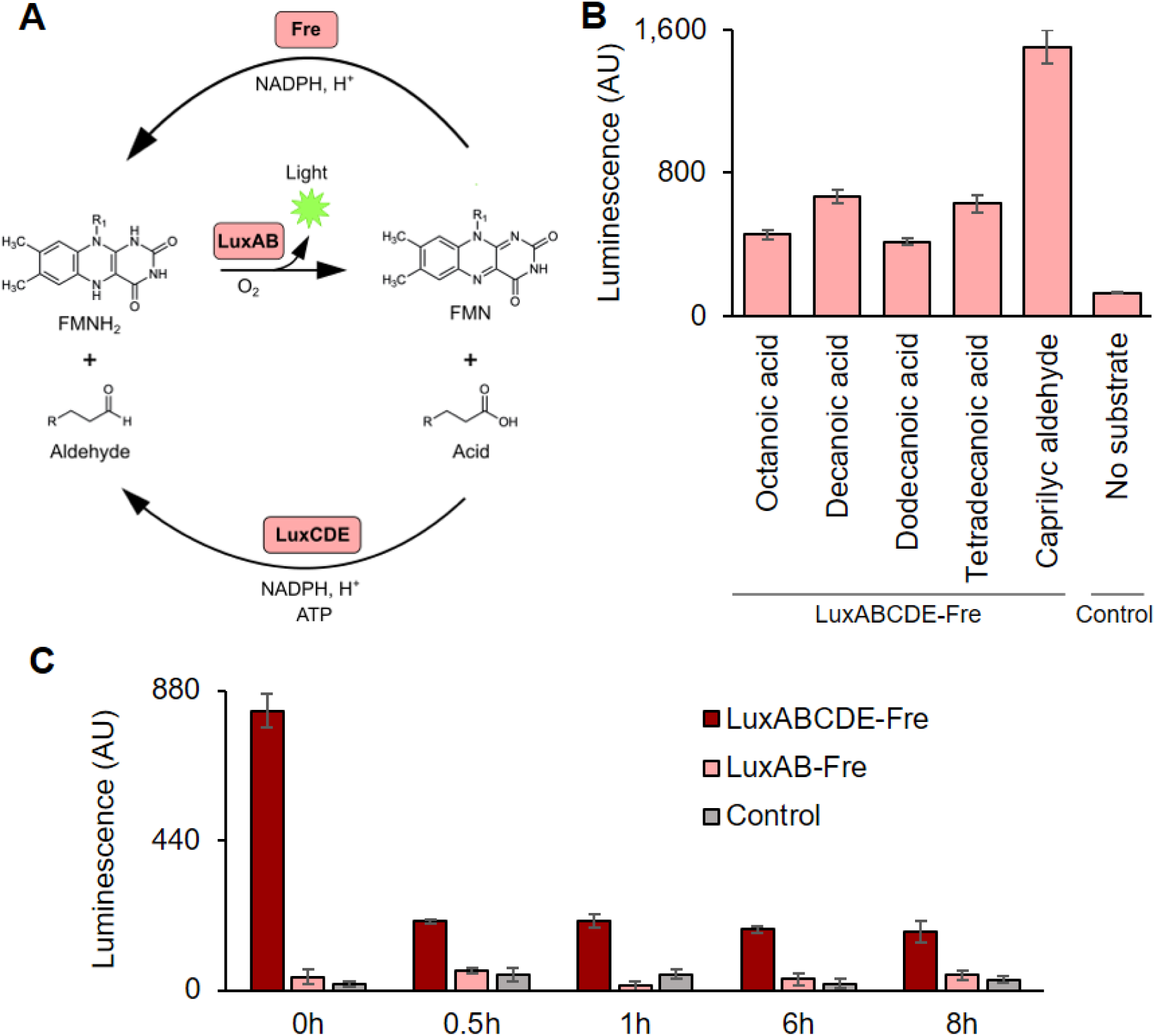
Substrate regeneration system with LuxABCDE-Fre. (A) The schematic of the LuxABCDE-Fre substrate regeneration system. LuxAB generates light with reduced flavin mononucleotide (FMNH_2_) and long-chain aldehyde; those substrates are converted into oxidized flavin mononucleotide (FMN) and corresponding long-chain acid. NAD(P)H-flavin reductase (Fre) reduces FMN back to FMNH_2_, and LuxCDE reduces the acid back to the corresponding aldehyde. (B) Luminescence measurement with the LuxABCDE-Fre system. LuxAB-Fre and LuxCDE were expressed in TXTL and mixed with 1 mM long-chain fatty acids (octanoic acid, decanoic acid, dodecanoic acid, or tetradecanoic acid) or caprylic aldehyde, followed by luminescence measurement. The reaction also contained FMN, NADPH, and ATP. Control represents a reaction using TXTL without enzyme expression. (C) Luminescence kinetics measurement with decanoic acid. LuxAB-Fre and LuxCDE were expressed in TXTL. For LuxABCDE-Fre reaction, the TXTL expressing LuxAB-Fre and LuxCDE were mixed with 1 mM decanoic acid (time = 0). For LuxAB-Fre reaction, the TXTL expressing LuxAB-Fre was used. For Control reaction, TXTL without enzyme expression was used. The reaction also contained FMN, NADPH, and ATP. The luminescence was measured after 0.5, 1, 6, and 8 hours. The graphs show means with error bars that signify SEM (n = 3).

### A simplified luciferase based expression analysis using HiBiT NanoLuc

Because the availability of resources limits the TXTL’s ability to express genes, reducing the metabolic burden for the reporter genes is highly advantageous. As a part of luciferase optimizations for TXTL, we built another burden-reduced TXTL with NanoLuc. NanoLuc can be split into two parts: LgBiT (18 kDa subunit derived from N-term NanoLuc) and HiBiT (1.3 kDa peptide, 11 amino acids, derived from C-term NanoLuc)^26,27^. To make a cell-free extract, we used the *E. coli* carrying a plasmid of the LgBiT gene, a bigger fragment. This LgBiT-carrying TXTL only requires 11 amino acids (HiBiT) expression for luminescence (**Fig. 5A**). We also confirmed that we could fuse a protein on either end of HiBiT, with eGFP fluorescence increased when it was fused with the C-term of HiBiT (**Fig. 5B**). Both N-term and C-term HiBiT fusion generated brighter signals than HiBiT without fusion proteins in LgBiT carrying TXTL (**Fig. 5C**). When we measured the kinetics of HiBiT expression, HiBiT with C-term eGFP fusion generated luminescence earlier (max at 10 minutes) than with N-term eGFP fusion (max at 40 minutes) **(Fig. 5D)**.

**Figure 5.**
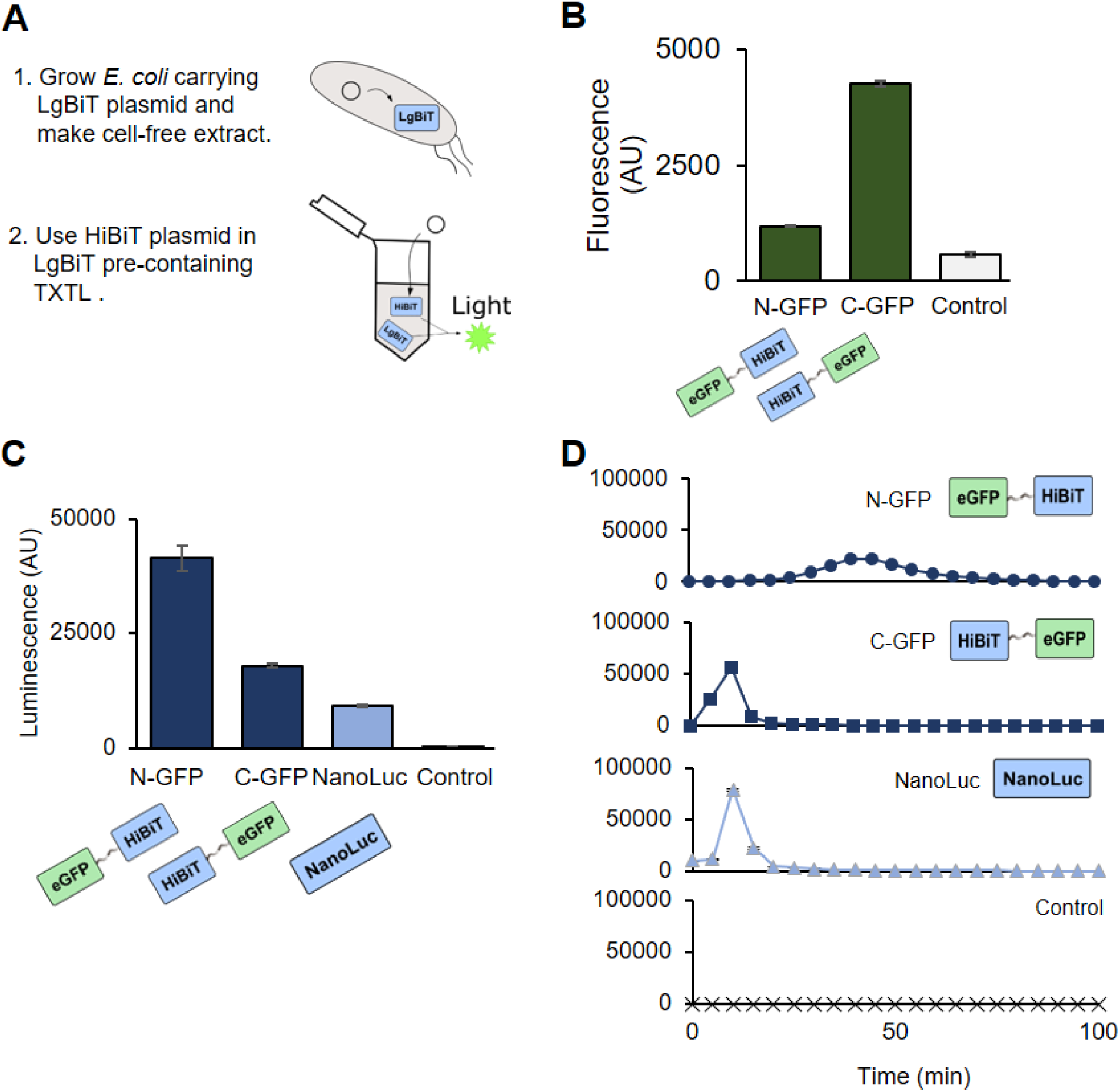
HiBiT reporter system. (A) The schematic of how LgBiT-carrying *E. coli* cell-free extract works. The plasmid coding LgBiT gene under the sigma 70 promoter is transformed into *E. coli* Rosetta 2 strain. The cell-free extract is made with that strain; thus, the extract contains LgBiT. Once HiBiT is expressed in TXTL, HiBiT produces luminescence by reconstituting full luciferase with LgBiT. (B) eGFP fluorescence measurement of HiBiT-GFP fusion proteins. Fusion proteins were expressed in TXTL at 30°C for 8 hours, followed by the measurement. (C) End-point luminescence assay with fusion HiBiTs. The fusion HiBiTs were expressed in LgBiT-containing TXTL at 30°C for 8 hours. After the expression, 1 µM Furimazine was added, followed by luminescence measurement. (D) Luminescence kinetics measurement with LgBiT-containing TXTL. HiBiT plasmids and 1 µM Furimazine were added at the TXTL reaction set up and incubated at 30°C. The luminescence was measured every 5 minutes during the TXTL reaction. N-GFP, N-terminal eGFP fusion with HiBiT; C-GFP, C-terminal eGFP fusion with HiBiT; Control, reaction without enzyme expression. The graphs show means with error bars that signify SEM (n = 3).

## Discussion

### Expression confirmation in the substrate-multiplexing assay

In the substrate-multiplexing assay (**Fig. 2C-F**), we confirmed that luciferases were expressed in the reactions by Western blot analysis (**Fig. S1**). The LuxAB-Fre reaction is not shown in **Fig. S1**, because His-tagged Fre did not show up on the membrane (LuxA and LuxB were not attached to His-tags). His-tagged Fre was detected on a membrane when expressed by itself (**Fig. S8**). We think the absence of the Fre band was because of the metabolic burden of expressing three genes; Fre’s reduced expression resulted in invisible bands on the membrane. Since we detected the luminescence, we consider the enzymes were still expressed enough to generate a measurable signal.

### Substrate regeneration with LuxABCDE-Fre

We tested the LuxABCDE-Fre substrate regeneration system with octanalydehyde (**Fig. S7**). The luminescence was the highest at 0.5 hours, while the starting time (t = 0) was the highest for the decanoic acid reaction (**Fig. 4C**). When preparing the reaction with octanalydehyde, we kept the samples on ice for 10 minutes while recovering the connection error between a plate reader and a computer. We think this cooled the reactions and caused the delay in the maximum luminescence reaction for the reaction in **Fig. S7**. The current limitation of the LuxABCDE-Fre substrate regeneration is that we need separately express LuxAB-Fre and LuxCDE in two TXTL reactions. We tried expressing LuxABCDE-Fre in a single TXTL reaction; however, we did not detect luminescence in that reaction. For the practical applications using the entire pathway, further optimization of TXTL conditions is required. We believe substrate regeneratively is still useful, particularly once we achieve the control of larger multiple gene networks in TXTL.

### Substrate regeneration with fungi luciferase

We tried to reconstitute the known substrate regeneration pathway of the fungi luciferase. After Luz produces light and caffeylpyruvic acid, CPH-Hisps-NPGA pathway brings caffeylpyruvic acid back to hispidin (**Fig. S10**)^18–20^. However, we could not reproduce those reactions in TXTL, probably because of unidentified co-factors or post-translational modifications. *E. coli* TXTL might not be suitable for post-translational modifications required for some, still unidentified, eukaryotic enzymes. See the supplemental information section, “Efforts to engineer the fungi luciferase substrate regeneration pathway.”

### Fusion proteins with luciferase reporters

In the eGFP fusion experiment, we connected eGFP and the luciferases with GS-linker (GGGGS) repeats. We chose eGFP because of the easiness of measurement and GS-linker because of the most used peptide linkers. We tried different lengths of GS liners (3x and 2x) but did not see the expression improvement (**Fig. 3D, Fig. S3, Fig. S4AB, Fig. S5, Fig. S6**, and **Fig. S9**). We failed in finding the constructs that generate both luminescence and fluorescence for LuxA and LuxB (**Fig. S4**). However, we think there is still a chance that LuxA and LuxB can be used as fusion constructs with further optimizations of the combination of linkers and fusion proteins. In **Fig. 5D**, the C-term eGFP fused HiBiT luminescence earlier than the N-term fusion construct. This faster luminescence is probably because the C-term fusion expresses HiBiT first and can complete the NanoLuc formation. In contrast, the N-term eGFP fusion construct cannot luminesce until the whole GFP-NanoLuc is expressed.

### Summary and perspectives

In this work, we established the use of two new luciferase systems as protein expression reporters in cell-free translation systems, and we demonstrated a technique that minimizes metabolic burden on TXTL while using split luciferase assay.

We characterized performance of those new luciferases in TXTL. Both Luz and LuxAB did not show cross-reactivity with commonly used luciferase substrates (**Fig. 2B-F**). Luz was still active when it fused with eGFP, showing potential to be used as a fusion protein (**Fig. 3CD**).

We successfully demonstrated a substrate regenerative luciferase reaction with LuxABCDE-Fre, enabling long time course kinetic experiments with luciferase readout in TXTL (**Fig. 4**).

In an effort to engineer a streamlined luciferase readout in TXTL, we also explored the use of small peptide tag HiBiT, which produces luciferase activity upon binding to larger domain to complete NanoLuc luciferase (**Fig. 3AB** and **Fig. 5AB**).^26,27^

Together, the work presented in this paper expands the toolbox of luminescent protein reporters for cell-free applications, building a more complete and versatile platform for a variety of cell-free and synthetic cell applications.

## Acknowledgments

We thank Dr. Arjun Khakhar for providing insights and information on the fungal luciferase pathway. We thank Dr. Daniel Voytas and Dr. Arjun Khakhar for the gift of the plasmid contains five enzymes for the fungi luciferase pathway. We thank Dr. Vincent Noireaux for the gift of T7 RNA polymerase plasmid. We thank Evan Kalb for the discussion of the HiBiT/LgBiT luciferase system.

This work was supported by NIH award 5R01MH114031 RNA Scaffolds for Cell Specific Multiplexed Neural Observation, NASA award 80NSSC18K1139 Center for the Origin of Life - Translation, Evolution And Mutualism, NSF award 1807461 SeMiSynBio Very Large scale genetic circuit design and automation, NSF award 1840301 RoL:FELS:RAISE Building and Modeling Synthetic Bacterial Cells, and the Funai Foundation for Information Technology.

## Materials and Methods

DNA oligonucleotides were purchased from Integrated DNA Technologies (IDT). Thermal cyclers used for sample incubation were a Bio-Rad T100 thermo cycler running software version 1.201. A plasmid for fungi luciferase pathway, P307-FBP_6, was a gift from Daniel Voytas lab at the University of Minnesota^20^. Cloning vector plasmids, pCI-T7Max-UTR1-CTerminus8xHis-T500 and pCI-T7Max-UTR1-NTerminus8xHis-T500, were obtained from our lab stock^25^. Plasmids for other luciferases, pGreen_dualluc_3’UTR_sensor, pGEN-luxCDABE, pUAS-NanoLuc, and pBad-LgBiT-PhoCl1-SmBiT-MBP, were purchased from Addgene^28– 30^.

### TxTl reactions

This protocol was adapted from Noireaux^4^ and Jewett^5^ protocols. The Rosetta 2 (Novagen, 71400) cell extract preparation was followed by the method described previously^31^ with one modification. A 750 ml 2xYPTG was grown at 30°C instead of 37°C. For H3H or LgBiT containing TXTL, Rosetta 2-derived strains carrying pLumi-H3H or pLumi-LgBiT were used for cell extract preparation. The electrocometent cells were prepared from Rosetta 2 *E. coli*, and the plasmid, pLumi-H3H or pLumi-LgBiT, was transformed. The successful transformant was selected through ampicillin resistance.

Cell-free transcription-translation (TXTL) reactions were composed of the following: 12 mM Magnesium glutamate; 140 mM potassium glutamate; 1 mM DTT; 1.5 μM T7 RNA polymerase; 0.4 U/µl Murine RNase Inhibitor (NEB, M0314S); 1x cell-free prep; 1x energy mix; and 1x amino acid mix. The plasmid concentrations were 15 nM. Unless otherwise specified, the TXTL reactions were incubated at 30°C for 8 hours, followed by 4°C hold.

10x Energy mix composition was the following: 500 mM HEPES, pH 8; 15 mM ATP; 15 mM GTP; 9 mM CTP; 9 mM UTP; 2 mg/mL E. coli tRNA; 0.68 mM Folinic Acid; 3.3 mM NAD; 2.6 mM Coenzyme-A; 15 mM Spermidine; 40 mM Sodium Oxalate; 7.5 mM cAMP; 300 mM 3-PGA.

10x amino acid mix was prepared by mixing 20 mM of the following amino acids: alanine, arginine, asparagine, aspartic acid, cysteine, glutamic acid, glutamine, glycine, histidine, isoleucine, leucine, lysine, methionine, phenylalanine, proline, serine, threonine, tryptophan, tyrosine, and valine. Those amino acids were dissolved in pH 6.5, 400 mM potassium hydroxide solution.

### Western blot analysis

The Western blot was performed with a method described previously^25^. The samples were fractionated on a 37.5:1 acrylamide:bis-acrylamide SDS-page gel at 100V in 800 ml 1x SDS running buffer (25mM Tris, 192mM Glycine, 3.5mM SDS). The gel percentage and fractionation time varied and are indicated on each figure.

### Luciferase assays

#### Substrate preparation

The chemicals used in the luciferase assay were as follows: FMN-Na (Alfa Aesar, J66949.09), NADPH (Cayman Chemical Company, 9000743), ATP (Larova GmbH, ATP_100ML), D-luciferin (Cayman Chemical Company, 25836), coelenterazine (Cayman Chemical Company, 16123), coelenterazine H (Promega, S2011), frimazine (Aoblous, AOB36539), hispidin (Cayman Chemical Company, 10012605), octanaldehyde (Fisher Scientific, O004425ML), decyl aldehyde (Fisher Scientific, AC154971000), dodecyl aldehyde (fisher scientific), octanoic acid (Fisher Scientific, O002725ML), decanoic acid (Fisher Scientific, AC167271000), dodecanoic acid (Fisher Scientific, S25377), tetradecanoic acid (Fisher Scientific, AAA1206730). D-luciferin, furimazine, and hispidin were dissolved in DMSO as 10 mM stocks. Coelenterazine and Coelenterazine H were dissolved in ethanol as 10 mM stocks. Long-chain fatty aldehydes and acids were dissolved in ethanol as 500 mM stocks. Dodecyl aldehyde was not soluble in 100% ethanol at the concentration of 500 mM but used the suspension by vortexing every time before taking out.

#### Luminescence measurement setting

The luminescence measurements were performed with SpectraMax Gemini EM Microplate Reader or SpectraMax Gemini Microplate Reader. The readings were performed by measuring the luminescence of all the wavelengths with readings “6” and photomultiplier tube (PMT) setting “medium”. 15 µl of samples were transferred to a 384 well white flat bottom assay plate (Corning®, 3705) and measured. For the kinetics measurements, the plate was sealed tightly to avoid evaporation.

#### End-point luciferase assay

Enzymes were expressed in 20 μl TXTL with each plasmids’ concentration of 15 nM at 30°C for 6 hours. Then, in 50 μl luciferase reactions, the 20 μl TXTL, substrates, and co-factors were mixed. The substrate concentrations were 1mM for aldehydes (octanaldehyde, decyl aldehyde, or dodecyl aldehyde) or 10 μM for other substrates (D-luciferin, coelenterazine H, Furimazine, and Hispidin). FLuc reaction contained 5 mM MgCl_2_ and 1 mM ATP; fungi luciferase (H3H-Luz) reaction contained 1 mM NADPH; LuxAB+Fre reaction contained 100 μM FMN, 1 mM NADPH, and 1 mM ATP. For multiplexing assay, luciferase reactions were prepared with “All” the substrates or “All minus one” substrate. The substrate concentrations were 1 mM for octanaldehyde and 10 μM of other substrates (D-luciferin, coelenterazine H, Furimazine, and Hispidin). The reaction also contained 1 mM MgCl_2_, 1 mM ATP, 1 mM NADPH, and 100 μM FMN.

Immediately after mixing the reaction, luminescence was measured by a plate reader. For the control, water was added instead of the components.

#### LuxCDABE-Fre substrate regeneration assay

For the end-point measurement, TXTL 1 and TXTL 2 were prepared separately. TXTL 1 contained 15 nM LuxA, LuxB, and Fre plasmids, and TXTL 2 contained 15 nM LuxC, LuxD, and LucE plasmids. After incubating TXTL at 30°C for 6 hours, luciferase reactions were prepared with TXTL 1 and 2. The 50 μl luciferase reactions contained 20 μl TXTL 1, 20 μl TXTL 2, 100 μM FMN, 1 mM NADPH, 1 mM ATP, and 1 mM substrates (octanoic acid, decanoic acid, dodecanoic acid, 1-tetradecanoic acid, or octanaldehyde.) Immediately after mixing the luciferase reaction, the luminescence was measured by a plate reader. For the control reaction, water was added into the TXTL instead of the plasmid. The reaction components for the kinetics measurement were the same as the end-point measurement. The reading was performed every 5 minutes for 8 hours.

#### eGFP fluorescence measurement

The fluorescence was measured at λ_ex_ 488 nm and λ_em_ 509 nm with plate reader PMT setting “medium” and 6 reads per well. All fluorescence measurements were performed on SpectraMax. For the endpoint measurement, 19 μl of TXTL reaction was transported into a 384 black bottom well plate to measure.

## Supplementary information

for manuscript

### Expanding luciferase reporter systems for cell-free protein expression

by

Wakana Sato, Melanie Rasmussen, Christopher Deich, Aaron E. Engelhart, Katarzyna P. Adamala

**Figure S1.**
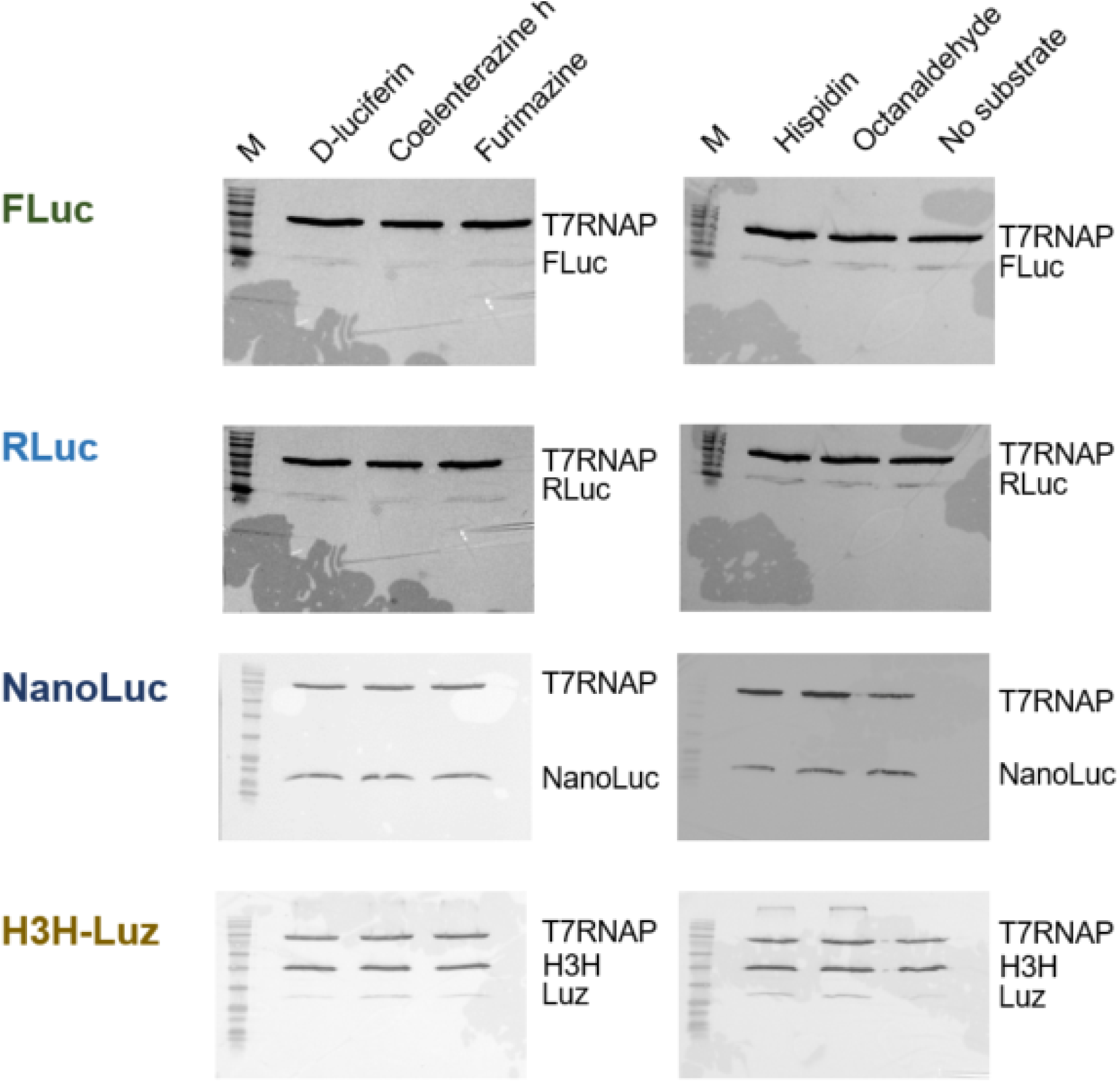
Western blot gels from the substrate specificity reactions. The loaded samples directly came from the reactions shown in **Fig. 2B**, indicating that the luciferase enzymes were expressed in the reactions. 15 μl of TxTL was loaded on each lane. FLuc and RLuc samples were fractionated on a 7.5% gel for 70 minutes at 100V. NanoLuc and H3H-Luz samples were fractionated on a 12% gel for 80 minutes at 100V. M, BLUEstain 2 Protein ladder, 5-245 kDa (Goldbio, P008-500); substrate names, a substrate that is contained in the reaction; T7RNAP, N-terminal His-tagged T7 RNA polymerase; FLuc, firefly luciferase with C-terminal His-tag; RLuc, Renilla luciferase with C-terminal His-tag; NanoLuc, NanoLuc luciferase with C-terminal His-Tag; H3H, hispidin-3-hydroxylase with C-terminal His-tag; Luz, fungi luciferase with C-terminal His-tag.

**Figure S2.**
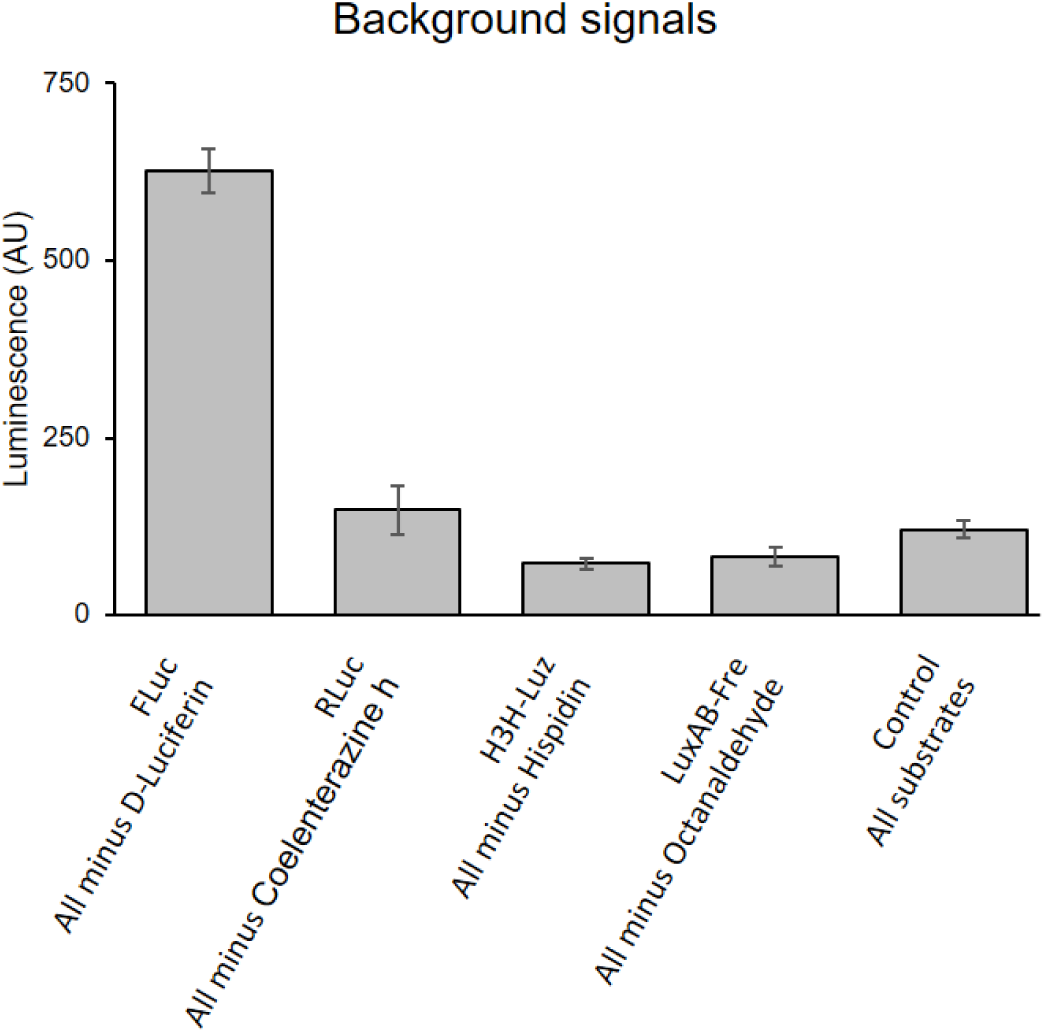
Background signals of the substrate specificity assay. The luciferases used in the reaction are indicated as FLuc, RLuc, H3H-Luz, or LuxAB-Fre. Control stands for reaction without enzyme expression. Substrates in the reaction were indicated as “All” (D-luciferin, coelenterazine h, hispidin, octanaldehyde) or “All minus one”, that one is the substrate supposed to react with the tested luciferase.

**Figure S3.**
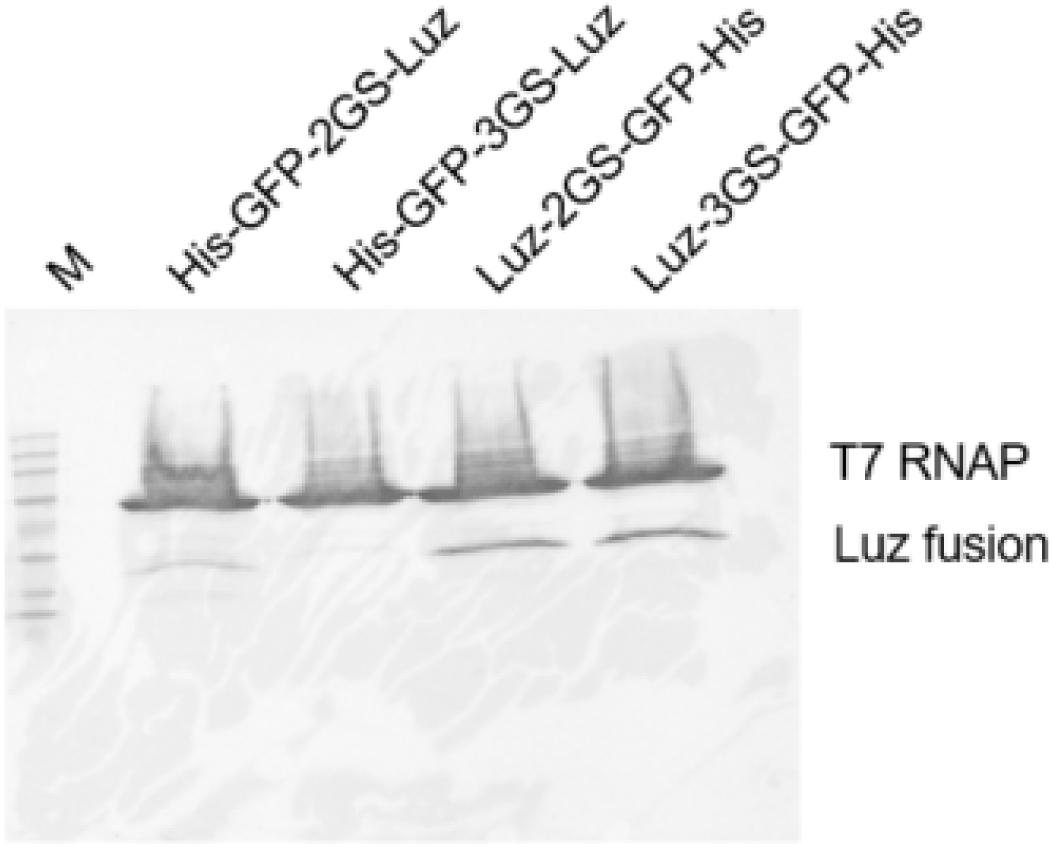
Luz-fusion gel image. Western blot of Luz-GFP fusion proteins. 15 μl of TxTL was loaded on each lane and fractionated on a 7.5% gel for 60 minutes at 100V. The lane labeled with M stands for BLUEstain 2 Protein ladder, 5-245 kDa (Goldbio, P008-500). The protein names used for lane labels represent the fusion protein constructs from N-terminal to C-terminal order. His, His-Tag; GFP, enhanced green fluorescent protein; 2GS, two GS-linker sequence (GGGGS) repeats; 3GS, three GS-linker sequence repeats; T7 RNAP, T7 RNA polymerase.

**Figure S4.**
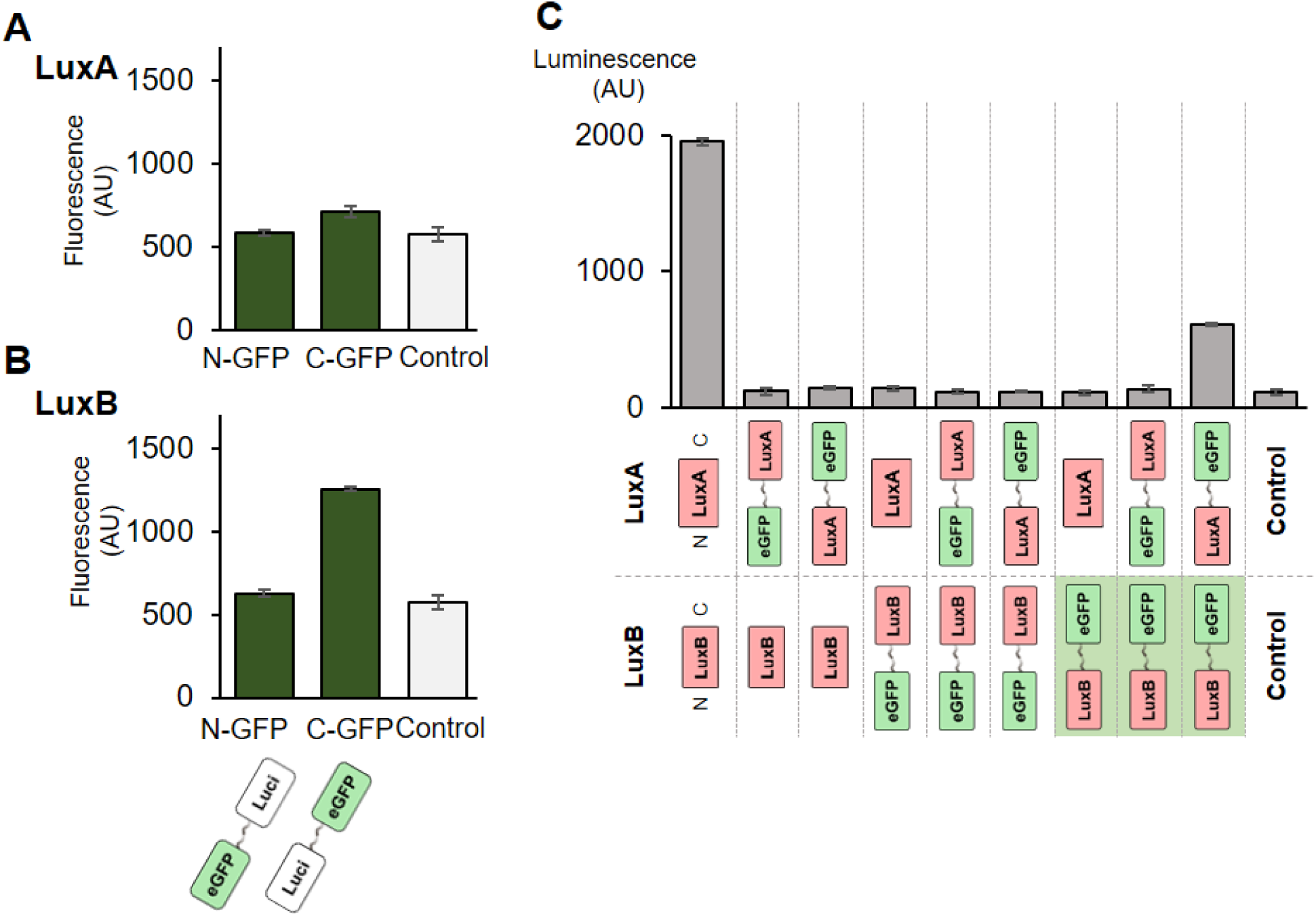
LuxA and LuxB capability as fusion proteins. (A and B) eGFP fluorescence measurement in TXTL. Fusion LuxA and LuxB were expressed in TXTL at 30°C for 8 hours, followed by the fluorescence measurement. (C) Luminescence measurement with all the combinations of LuxA and LuxB fusion constructs. LuxA and LuxB were expressed in TXTL, and then 1 mM Octanaldehyde was added, followed by luminescence measurement. Green shading behind the construct images indicates that the constructs fluorescence when expressed in TXTL. N-GFP, N-terminal eGFP fusion luciferase; C-GFP, C-terminal eGFP fusion luciferase; Control, reaction without enzyme expression.

**Figure S5.**
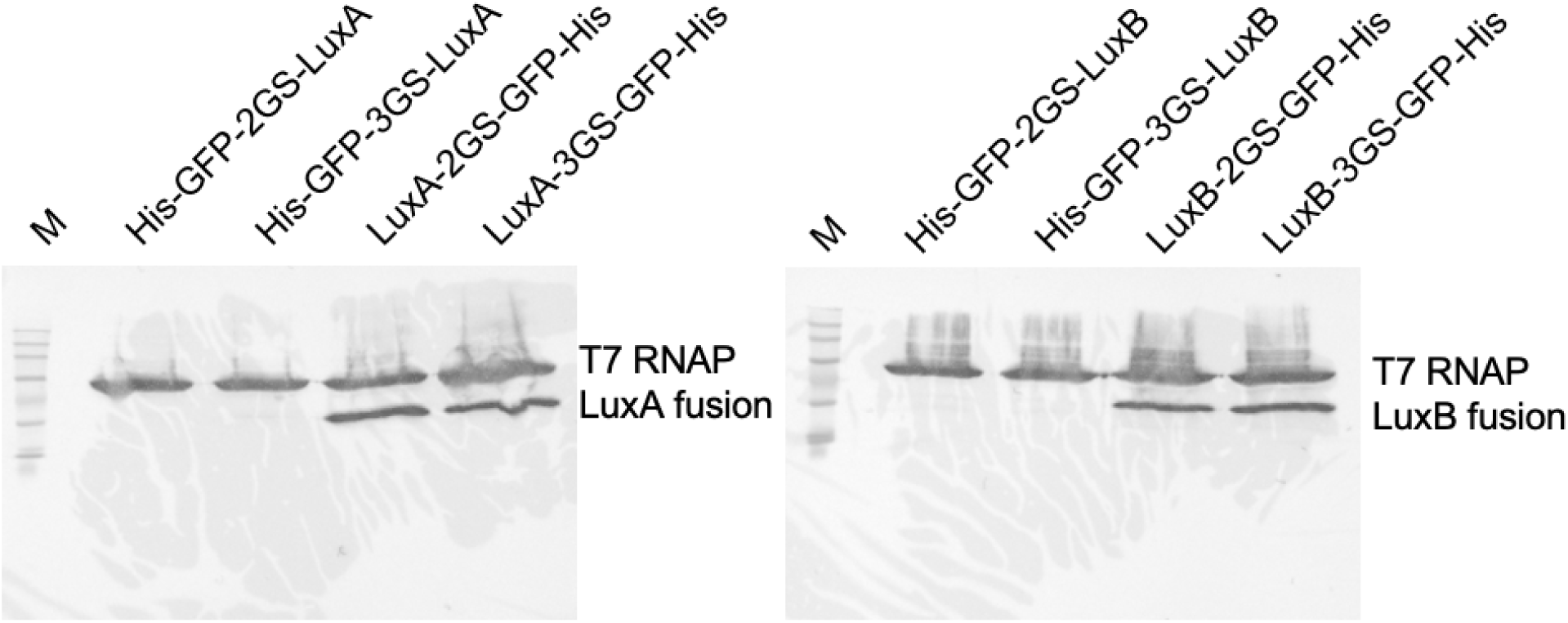
Western blot of LuxA- and LuxB-fusion proteins. 15 μl of TXTL was loaded on each lane and fractionated on a 7.5% gel for 60 minutes at 100V. The lane labeled with M stands for BLUEstain 2 Protein ladder, 5-245 kDa (Goldbio, P008-500). The protein names used for lane labels represent the fusion protein constructs from N-terminal to C-terminal order. His, His-Tag; GFP, enhanced green fluorescent protein; 2GS, two GS-linker sequence (GGGGS) repeats; 3GS, three GS-linker sequence repeats; T7 RNAP, T7 RNA polymerase.

**Figure S6.**
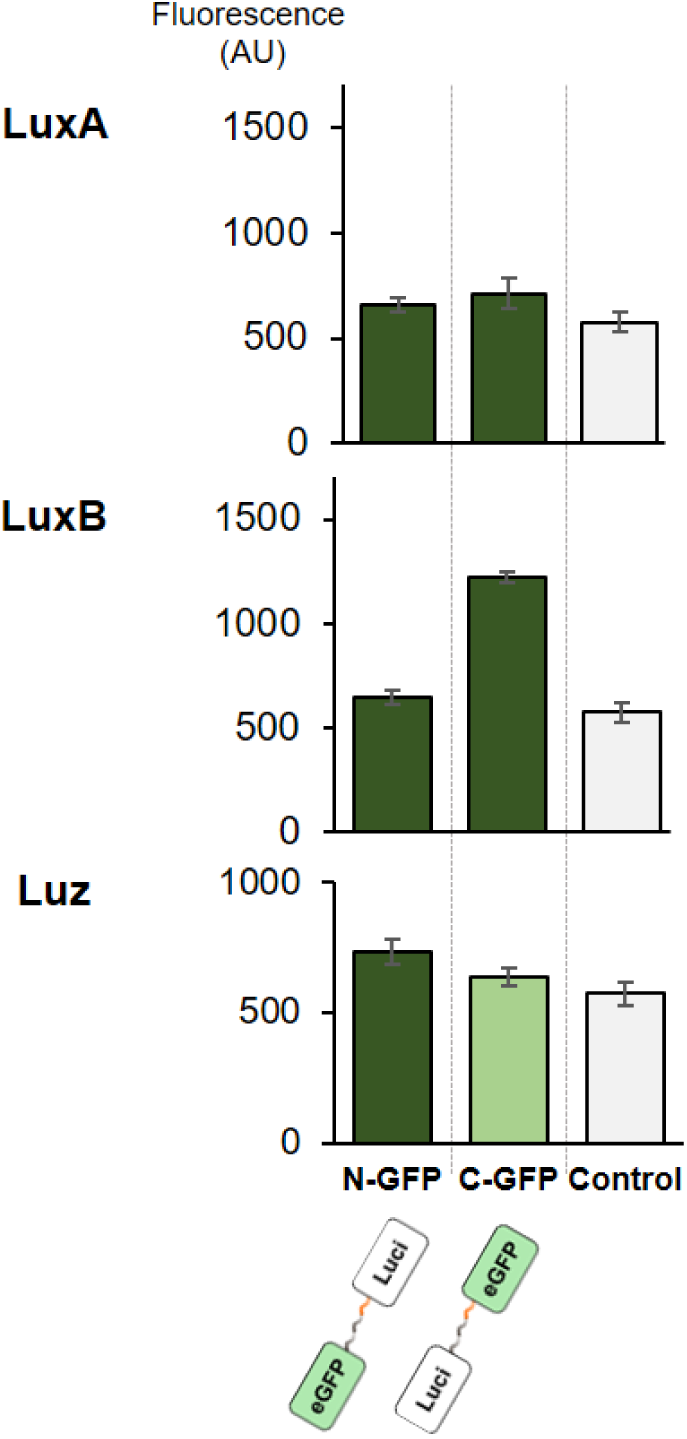
Fluorescence generated from eGFP-luciferase fusion proteins with extended GS-linker. All fusion proteins were expressed in TXTL at 30 °C for 8 hours, followed by fluorescence measurement. 19 μl of TXTL was used for the measurement. eGFP and luciferases are linked through 3x GS-linker (GGGGS.) Control stands for a reaction without protein expression.

**Figure S7.**
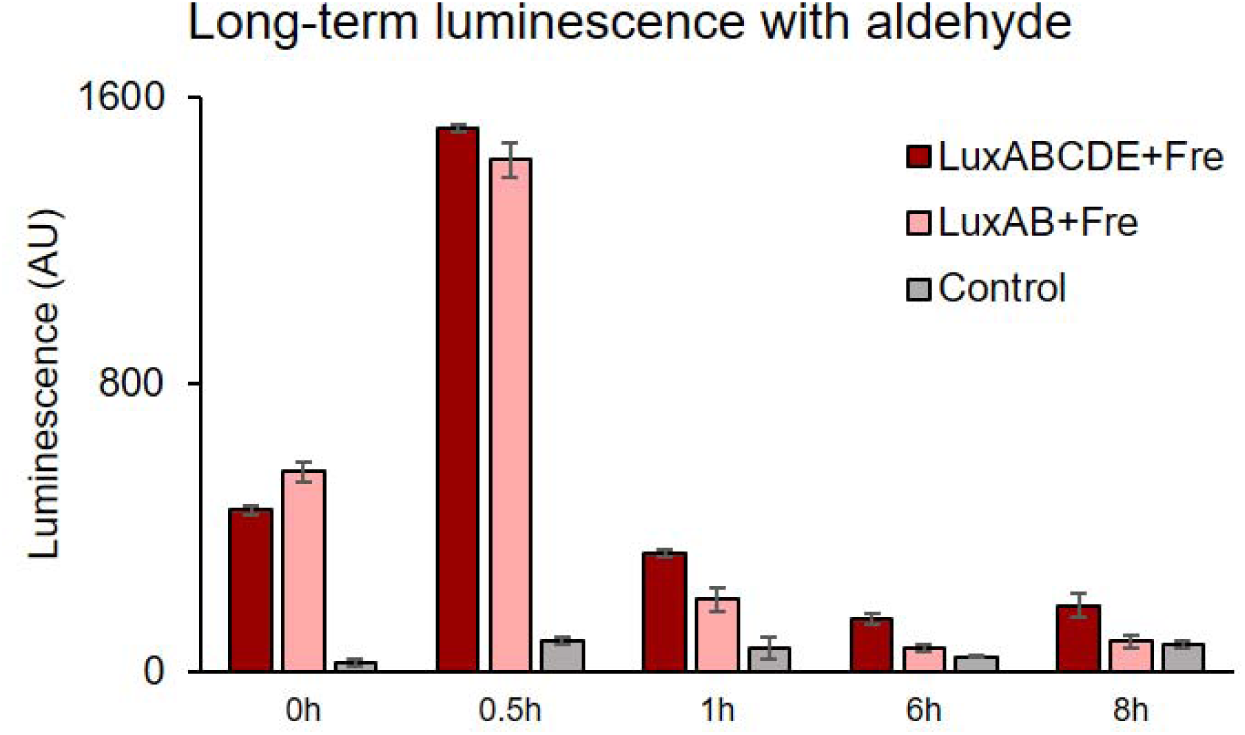
Luminescence kinetics measurement with octanaldehyde. After expressing LuxABCDE+Fre or LuxAB+Fre in TXTL, 1 mM octanaldehyde was added as the substrate (time = 0). The luminescence was measured after 0.5, 1, 6, 8 hours. LuxABCDE-Fre, a reaction with TXTL expressing LuxAB-Fre and LuxCDE; LuxAB-Fre, a reaction with TXTL expressing LuxAB-Fre; Control, reaction with TXTL without enzyme expression.

**Figure S8.**
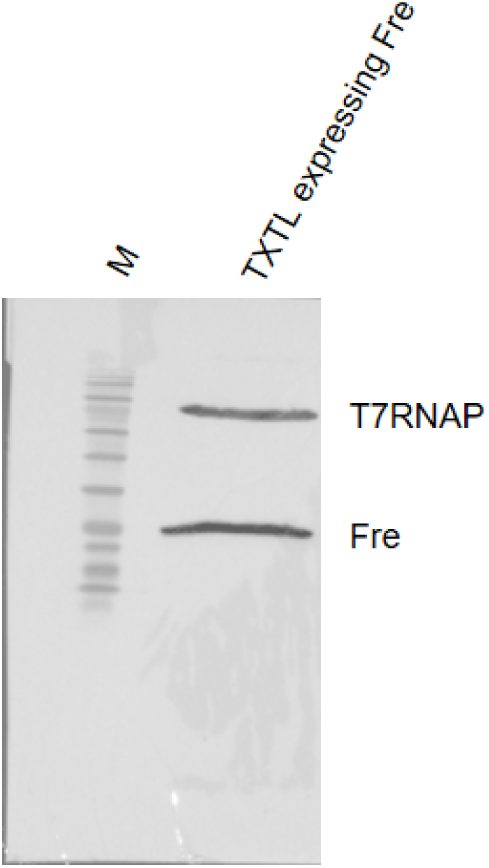
Western blot of Fre. 10 μl of TXTL expressing Fre was loaded on the right lane and fractionated on a 12% gel for 80 minutes at 100V. The left lane labeled with M stands for BLUEstain 2 Protein ladder, 5-245 kDa (Goldbio, P008-500).

**Figure S9.**
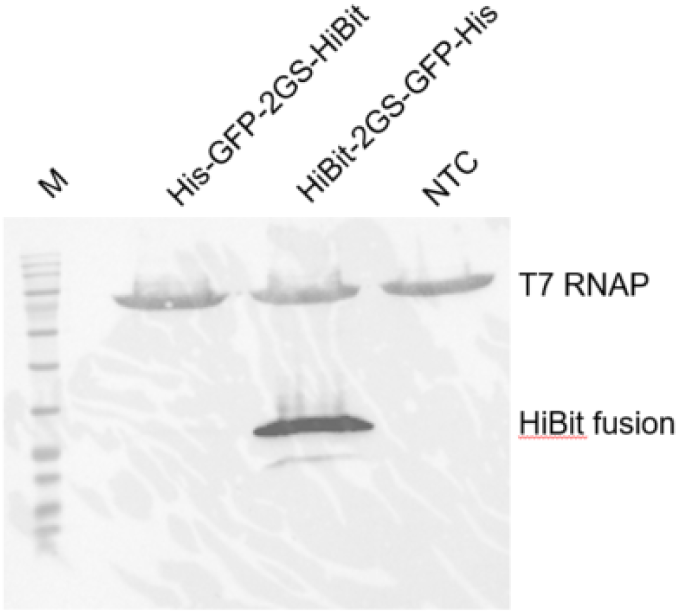
Western blot of HiBiT-fusion proteins. 15 μl of TXTL was loaded on each lane and fractionated on 7.5% gel for 60 minutes at 100V. The lane labeled with M stands for BLUEstain 2 Protein ladder, 5-245 kDa (Goldbio, P008-500). The protein names used for lane labels represent the protein structure from N-terminal to C-terminal order. His, His-Tag; GFP, enhanced green fluorescent protein; 2GS, two GS-linker sequence (GGGGS) repeats; T7 RNAP, T7 RNA polymerase.

**Figure S10.**
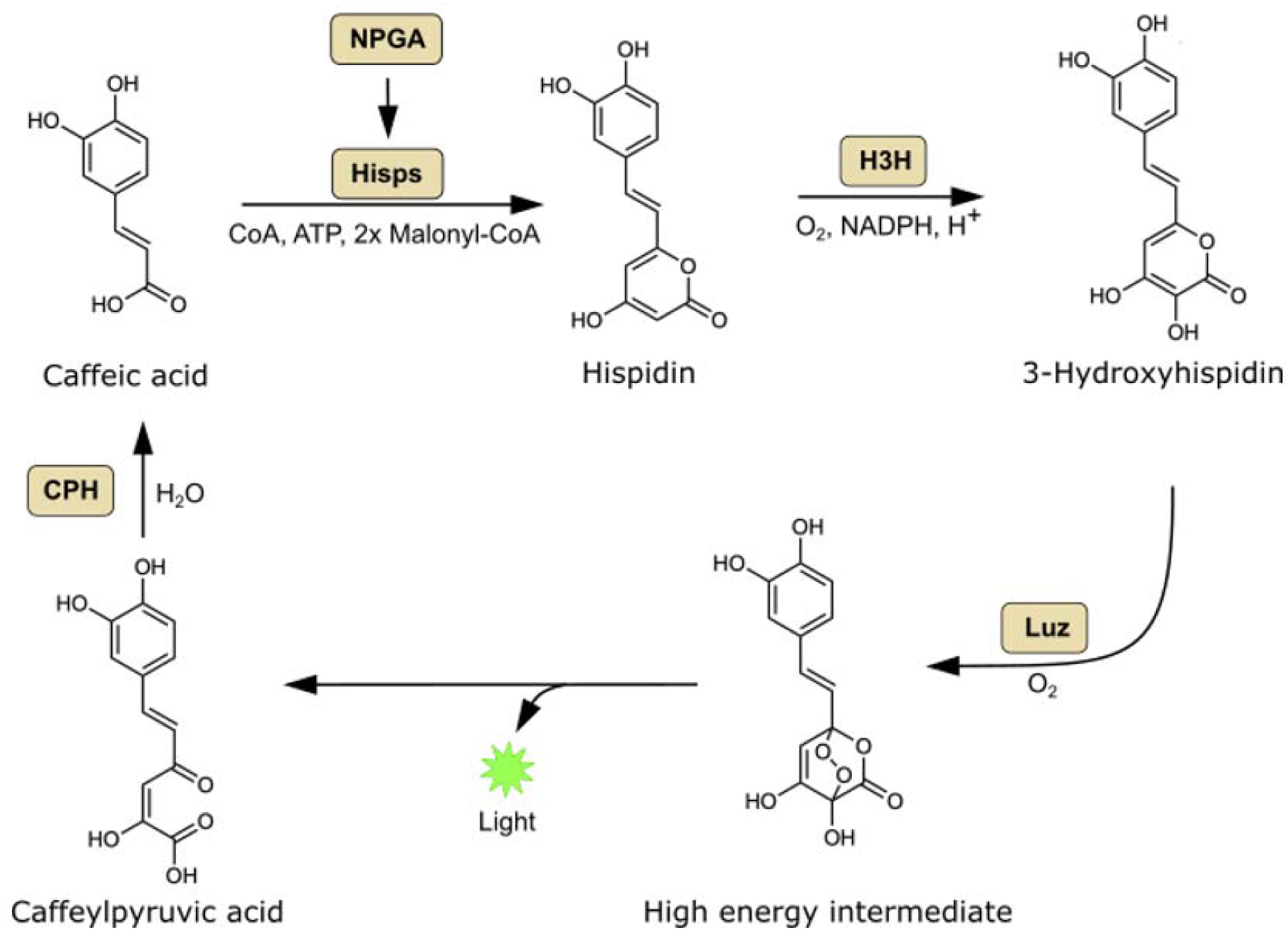
The proposed fungi luciferase substrate regeneration pathway. This figure was adapted from Khakhar et al. paper^20^. In the fungi luciferase system, hispidin is converted to 3-hydroxyhispidin by *Neonothopanus nambi* (*N. nambi*) hispidin-3-hydroxylase (H3H), and then *N. nambi* luciferase (Luz) yields light by reacting with 3-hydroxyhispidin. Caffeylpyruvic acid can be recycled into caffeic acid by caffeylpyruvate hydrolase (CPH). Caffeic acid can be converted into hispidin by hispidin synthase (Hisps). Hisps needs to be post-translationally activated by 4’-phosphopantetheinyl transferase (NPGA)^18–20^.

**Figure S11.**
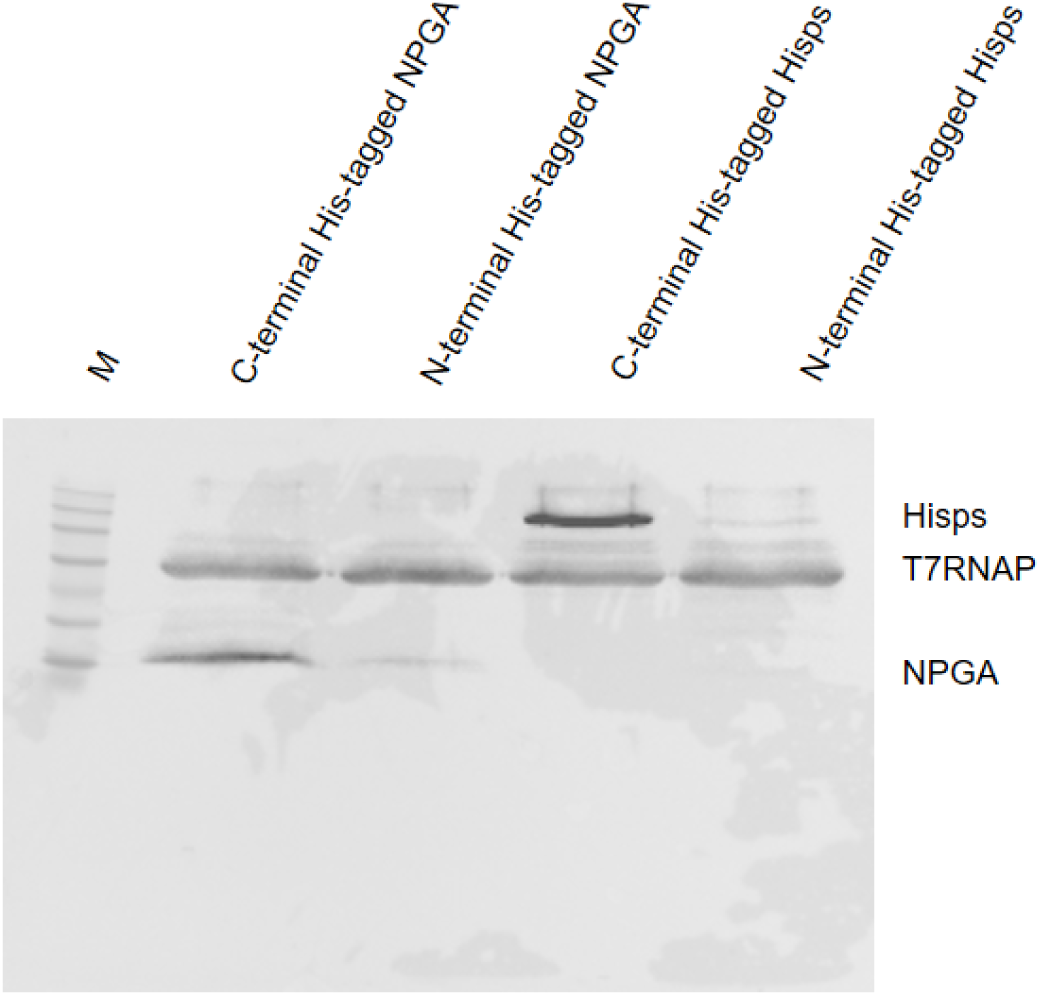
10 μl of TXTL was loaded on each lane and fractionated on 7.5% gel for 70 minutes at 100V. The left lane labeled with M stands for BLUEstain 2 Protein ladder, 5-245 kDa (Goldbio, P008-500). The other lanes were labeled with the protein names that were expressed in the TXTL. NPGA, 4’-phosphopantetheinyl transferase; Hisps, hispidin synthase; T7 RNAP, T7 RNA polymerase.

**Figure S12.**
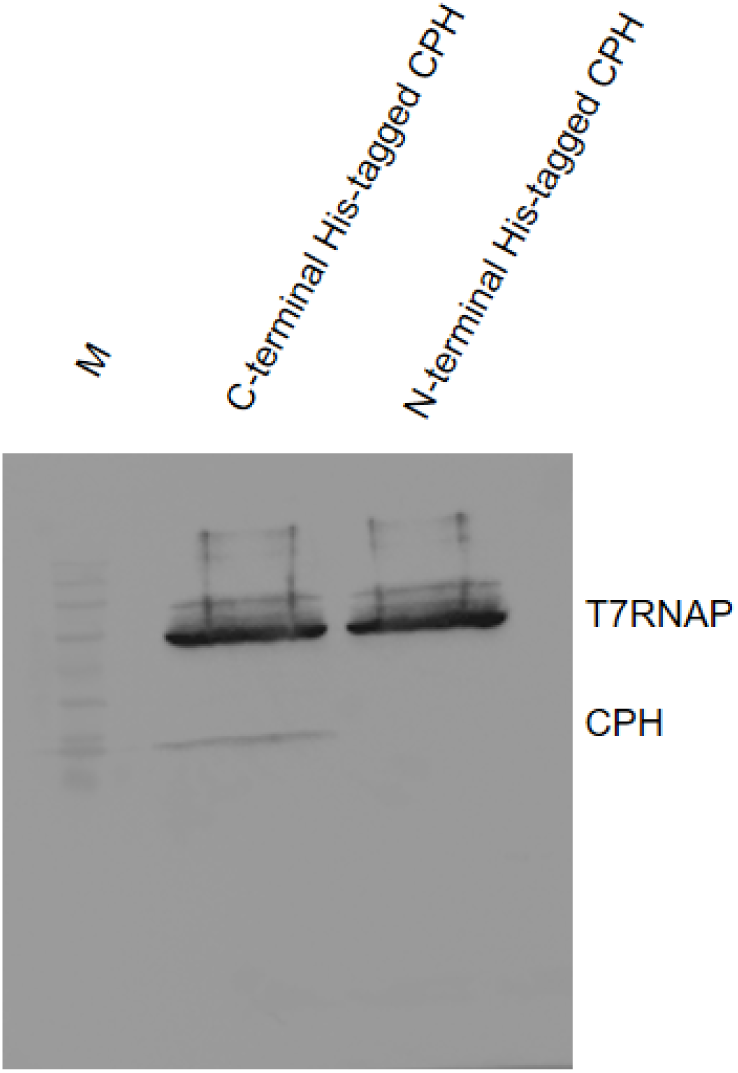
10 μl of TXTL was loaded on each lane and fractionated on 7.5% gel for 70 minutes at 100V. The left lane labeled with M stands for BLUEstain 2 Protein ladder, 5-245 kDa (Goldbio, P008-500). The other lanes were labeled with the protein names that were expressed in the TXTL. CPH, caffeylpyruvate hydrolase; T7 RNAP, T7 RNA polymerase.

**Figure S13.**
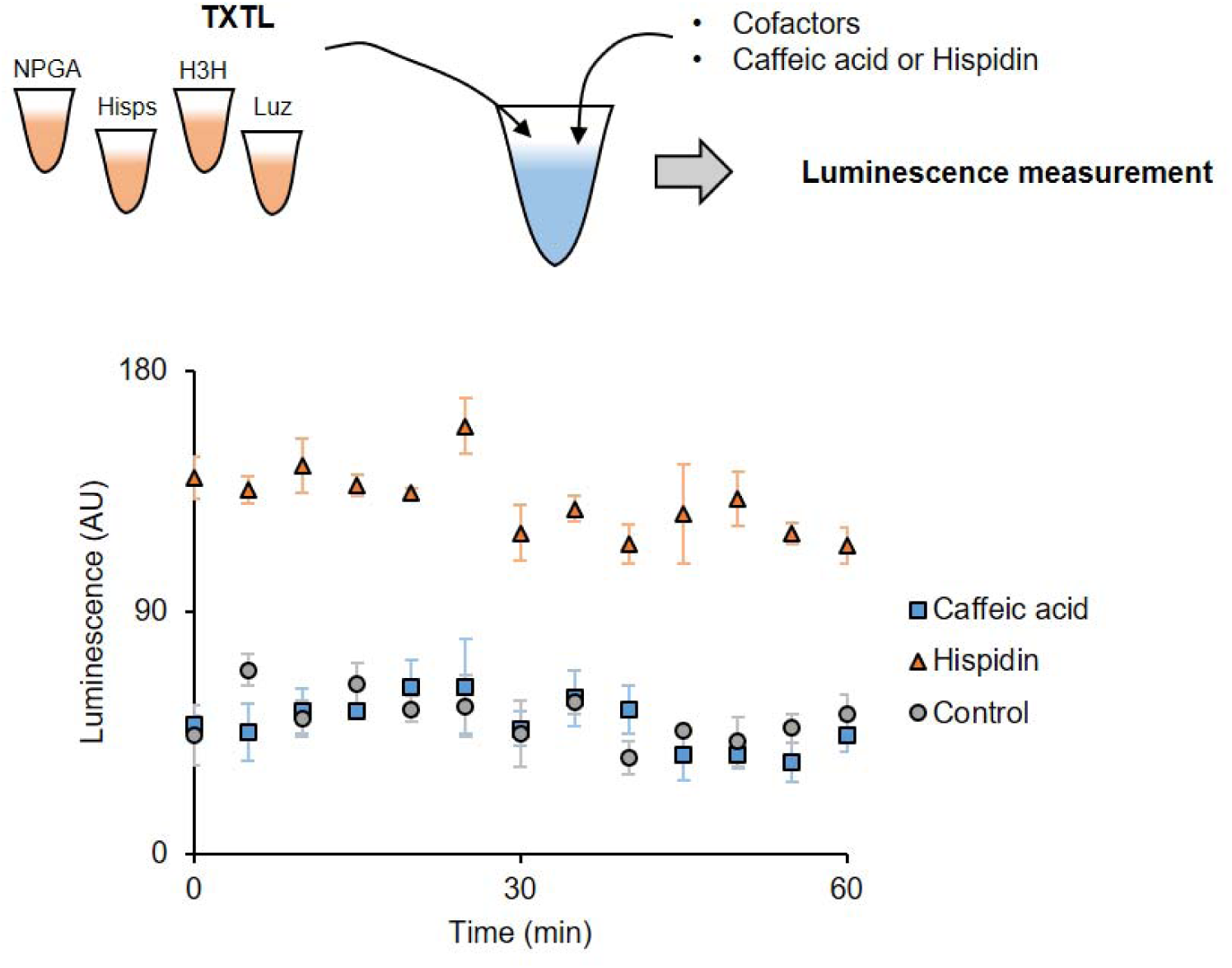
The result of caffeic acid conversion assay 1. NPGA, Hisps, H3H, and Luz were individually expressed in TXTL and mixed with cofactors and substrates (caffeic acid or hispidin). The luminescence was measured at 25°C for 1 hour, every 5 minutes. While the reaction with hispidin generated light, the reaction with caffeic acid did not. This indicates that the caffeic acid was not converted into hispidin. Control contains TXTLs without enzyme expression.

**Figure S14.**
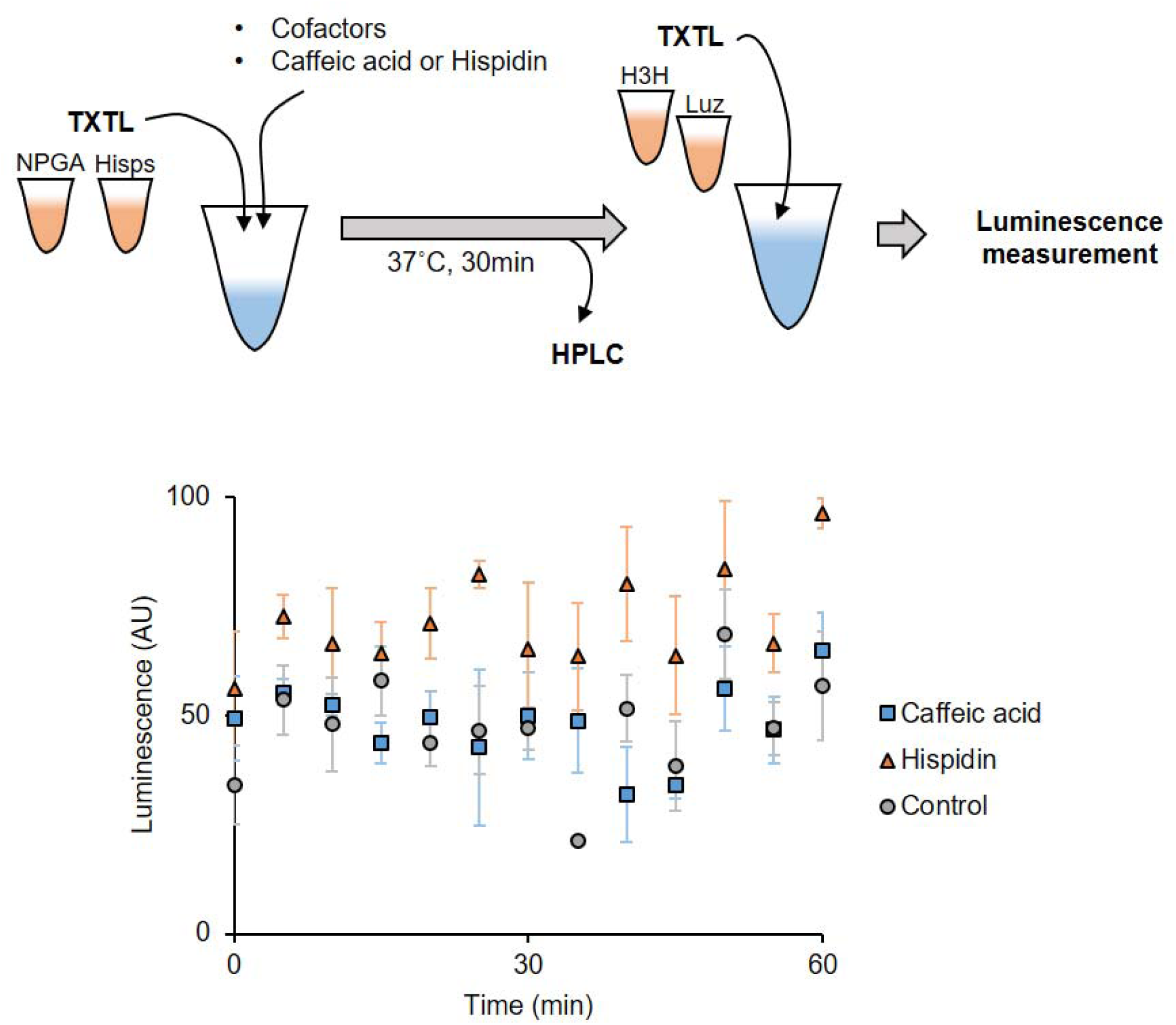
The result of caffeic acid conversion assay 2. NPGA, Hisps, H3H, and Luz were individually expressed in TXTL. NPGA and Hisps TXTLs were first mixed with cofactors and substrates (caffeic acid or hispidin). The reaction was incubated at 37°C for 30 minutes to facilitate phosphopanthetheinylation. The aliquot of the reaction was taken for HPLC analysis after this incubation. Then, H3H and Luz TXTLs were added to the reaction and measured the luminescence at 25°C for 1 hour, every 5 minutes. The reaction with hispidin slightly generated light. The reaction with caffeic acid was as same as Control reaction. This indicates that the caffeic acid was not converted into hispidin. Control stands for a reaction contains TXTLs without enzyme expression.

### Cloning methods

Primer sequences are listed in **Table S1**. Gene sequences are listed in **Table S2**. Plasmid names constructed in this paper are summarized in **Table S3**. Plasmids named with pTXTL contain a carbenicillin resistance gene and plasmids named with pLumi contain an ampicillin resistance gene for antibiotic selection. All the cloned plasmid sequences were verified by sequencing.

pTXTL-T7max-H3H, pTXTL-T7max-Luz, pTXTL-T7max-LuxA, pTXTL-T7max-LuxB, pTXTL-T7max-LuxC, pTXTL-T7max-LuxD, pTXTL-T7max-LuxE, pTXTL-T7max-Fre, pTXTL-T7max-FLuc, pTXTL-T7max-RLuc, and pTXTL-T7max-NanoLuc cloning.

The vector plasmid, pCI-T7Max-UTR1-CTerminus8xHis-T500, was digested with restriction enzymes at 37°C for 1 hour in 1x CutSmart Buffer (NEB, B7204). SpeI-HF (NEB, R3133L) and MluI-HF (NEB, R3198L) were used for H3H, Luz, Fre, FLuc, RLuc, and NanoLuc cloning. SpeI-HF and XhoI (NEB, R0146S) were used for LuxA, LuxB, LuxC, LuxD, and LuxE cloning. The digested plasmids were agarose gel purified using GenCatch Gel Extraction Kit (Epoch life science, 2260250). The purified vectors were treated with shrimp alkaline phosphatase (NEB, M0371L) at 37°C for 30 minutes, followed by the deactivation at 65°C for 15 minutes. Individual genes were amplified by Q5 High-Fidelity DNA Polymerase Master Mix (NEB, M0492L). H3H was amplified with P307-FBP_6, Primer1, and Primer2. Luz gene was amplified with gblock1, Primer3, and Primer4. Fre gene was amplified with gblock2, Primer15, and Primer16. FLuc and RLuc genes were amplified with pGreen_dualluc_3’UTR_sensor and primers (FLuc primers: Primer17 and Primer18, RLuc primers: Primer19 and Primer20). NanoLuc gene was amplified with pUAS-NanoLuc, Primer21, and Primer22. LuxA, LucB, LuxC, LuxD, and LuxB genes were amplified with pGEN-luxCDABE and primers (LuxA primers: Primer5 and Primer6, LuxB primers: Primer7 and Primer8, LuxC primers: Primer9 and Primer10, LuxD primers: Primer11 and Primer12, and LuxE primers: Primer13 and Primer14).

The PCR amplified genes were gel purified with GenCatch Gel Extraction Kit followed by restriction enzyme digestion, which the enzymes were the same as those that were used for vector digestion. The digested products were purified with GenCatch PCR Cleanup Kit (Epoch life science, 2360250). The vector and genes were ligated by T4 DNA Ligase (NEB, M0202L) in 1x T4 DNA Ligase reaction buffer (NEB, B0202S) at 16°C overnight. The following day, the ligase was deactivated at 65°C for 15 minutes and then transformed into *E. coli*.

#### pTXTL-T7max-HisGFPLuz, pTXTL-T7max-LuzGFPHis, and pLumi-T7max-H3H cloning

PCR reactions with Q5 High-Fidelity DNA Polymerase Master Mix were prepared to produce PCR fragments. For pTXTL-T7max-HisGFPLuz, (1) N-term-His-eGFP sequence was amplified with Primer23 and Primer24, (2) Luz sequence was amplified with Primer25 and Primer26, and (3) the backbone sequence, pCI-T7Max-UTR1-NTerminus8xHis-T500, was amplified with Primer27 and Primer28. For pTXTL-T7max-LuzGFPHis, (1) C-term-His-eGFP sequence was amplified with Primer29 and Primer30, and (2) Luz sequence was amplified with Primer31 and Primer32. For pLumi-T7max-H3H, (1) the backbone sequence, pGEN-luxCDABE, was amplified with Primer33 and Primer34, and (2) H3H was amplified with Primer35 and Primer36. The PCR products were purified with GenCatch PCR Cleanup Kit. The assembly was performed by circular polymerase extension cloning (CPEC) reaction, which was previously described^32^, then transformed into *E. coli*.

#### pTXTL-T7max-HisGFPHiBiT, pTXTL-T7max-HiBiTGFPHis, and pLumi-T7max-LgBiT cloning

PCR reactions with 1x Q5 High-Fidelity DNA Polymerase Master Mix were prepared to produce PCR fragments. For pTXTL-T7max-HisGFPHiBiT, (1) pTXTL-T7max-HisGFPLuz sequence was amplified with Primer37 and Primer38, and (2) pTXTL-T7max-HisGFPLuz was amplified with Primer39 and Primer40. For pTXTL-T7max-HiBiTGFPHis, (1) pTXTL-T7max-LuzGFPHis was amplified with Primer41 and Primer37, and (2) pTXTL-T7max-LuzGFPHis was amplified with Primer42 and Primer40. For pLumi-T7max-LgBiT, (1) pBad-LgBiT-PhoCl1-SmBiT-MBP was amplified with Primer43 and Primer44, and (2) pGEN-luxCDABE was amplified with Priemr45 and Primer33, and (3) pGEN-luxCDABE was amplified with Primer46 and Primer34. The PCR products were purified with GenCatch PCR Cleanup Kit. The assembly was performed with Gibson Assembly 1x Master Mix (NEB, E2611L) and incubated at 50°C for 30 minutes. The assembled products were transformed into *E. coli*.

### Efforts to engineer the fungi luciferase substrate regeneration pathway

NPGA, Hisps, and CPH are required in addition to Luz-H3H system for reconstructing the fungi substrate regeneration pathway (**Fig. S10**). First, we cloned NPGA, Hisps, and CPH genes into a vector plasmid, pCI-T7Max-UTR1-CTerminus8xHis-T500 or pCI-T7Max-UTR1-NTerminus8xHis-T500^25^. Those genes’ expressions were confirmed by Western blot (**Fig. S11, S12**). We used C-terminal His-tag constructs for NPGA, Hisps, and CPH, in later experiments, because of more robust expression in TXTL (**Fig. S11, S12**).

First, we expressed NPGA, Hisps, H3H, and Luz individually in 4 different TXTLs at 30°C for 8 hours. Then, in 90 µl reaction, we mixed the following components: 20 µl of each TXTLs, 2 mM ATP, 2 mM malonyl CoA, 1 mM CoA, and 1 mM NADPH. For substrate, we added 2 mM Caffeic acid or 200 µM Hispidin. We incubated the reaction at 25°C and measured the luminescence for 1 hour, every 5 minutes. However, only Hispidin-containing reaction produced luminescence (**Fig. S13**).

In a previous report, *in vitro* phosphopanthetheinylation assay with NPGA was performed at 37°C, instead of 30°C (our TXTL reaction)^33^. Thus, we split the reaction into two steps: (1) Conversion of caffeic acid to hispidin and (2) light generative reaction from hispidin. Briefly, we expressed NPGA, Hisps, H3H, and Luz individually at 30°C for 8 hours. Then, we mixed NPGA and Hisps TXTLs with 2 mM ATP, 2 mM malonyl CoA, 1 mM CoA, 1 mM NADPH, and 10 mM MgCl_2_. MgCl_2_ was added to make the reaction condition similar to the previous *in vitro* assay^33^. The reaction was incubated at 37°C for 30 minutes. The aliquot of samples was analyzed by high-performance liquid chromatography (HPLC); however, we did not detect the conversion from caffeic acid to hispidin. We then added H3H and Luz TXTLs to the reaction mixture and measured the luminescence at 25°C for 1 hour, every 5 minutes. We observed slight luminescence from a reaction containing hispidin as the substrate; however, the reaction containing caffeic acid did not generate any light (**Fig. S14**).

Lastly, we purified both NPGA and Hisps through His-tagged protein purification and performed an *in vitro* assay to see the conversion from caffeic acid to hispidin. The 40 µl reaction contained 100 mM Tris-HCl buffer (pH6.8), 1 µM NPGA, 1 µM Hisps, 2 mM ATP, 2 mM malonyl CoA, 500 µM CoA, and 40 mM MgCl_2_. The substrate was either 2 mM caffeic acid or 200 µM hispidin. The reaction was incubated at 37°C for 30 minutes. After the incubation, we stopped the reaction by adding 200 µl of ethanol, and the liquid was evaporated completely by a speed vacuum concentrator. The pellets were dissolved in methanol to inject HPLC. However, we could not detect the conversion of caffeic acid to hispidin by HPLC.

**Table S1.**
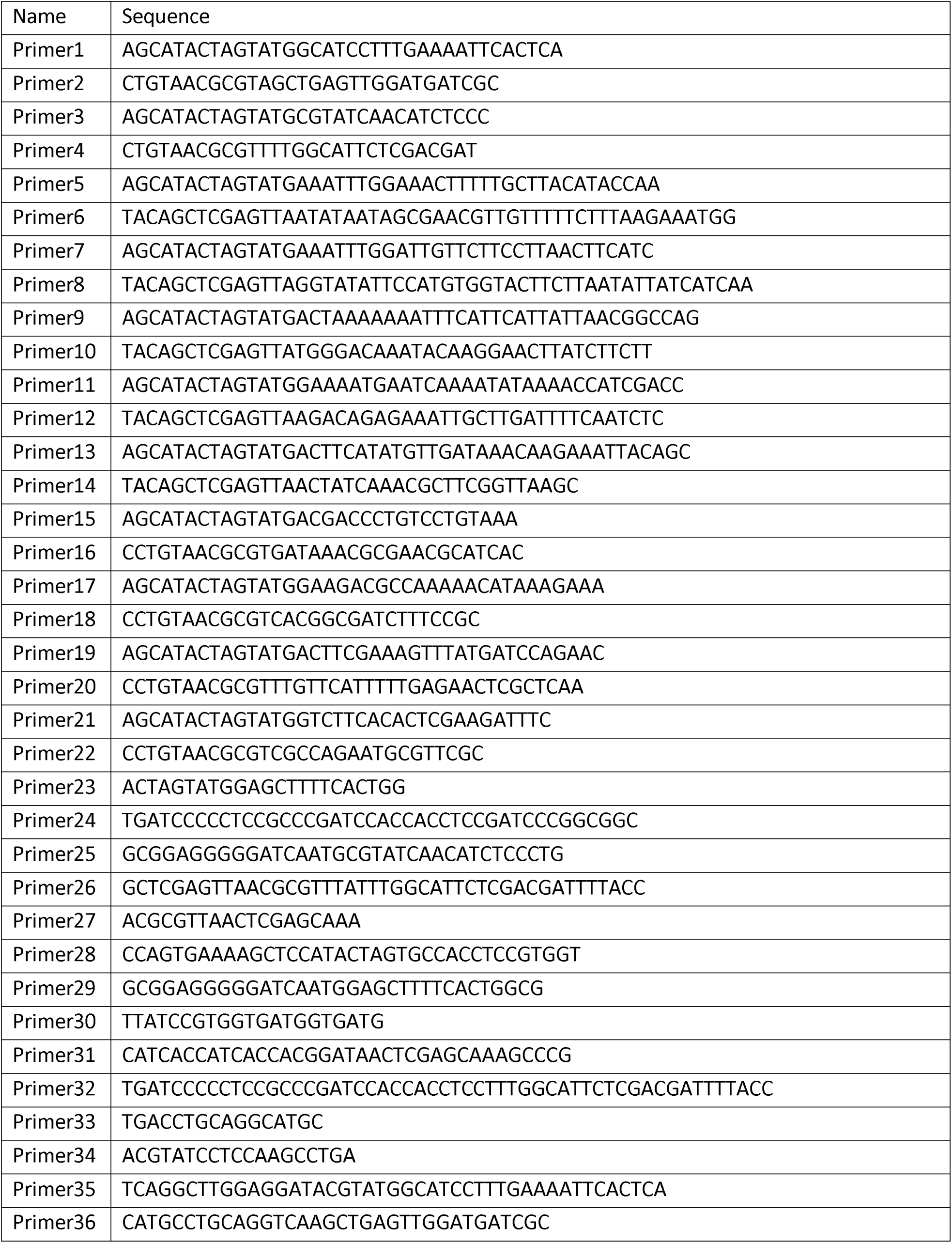
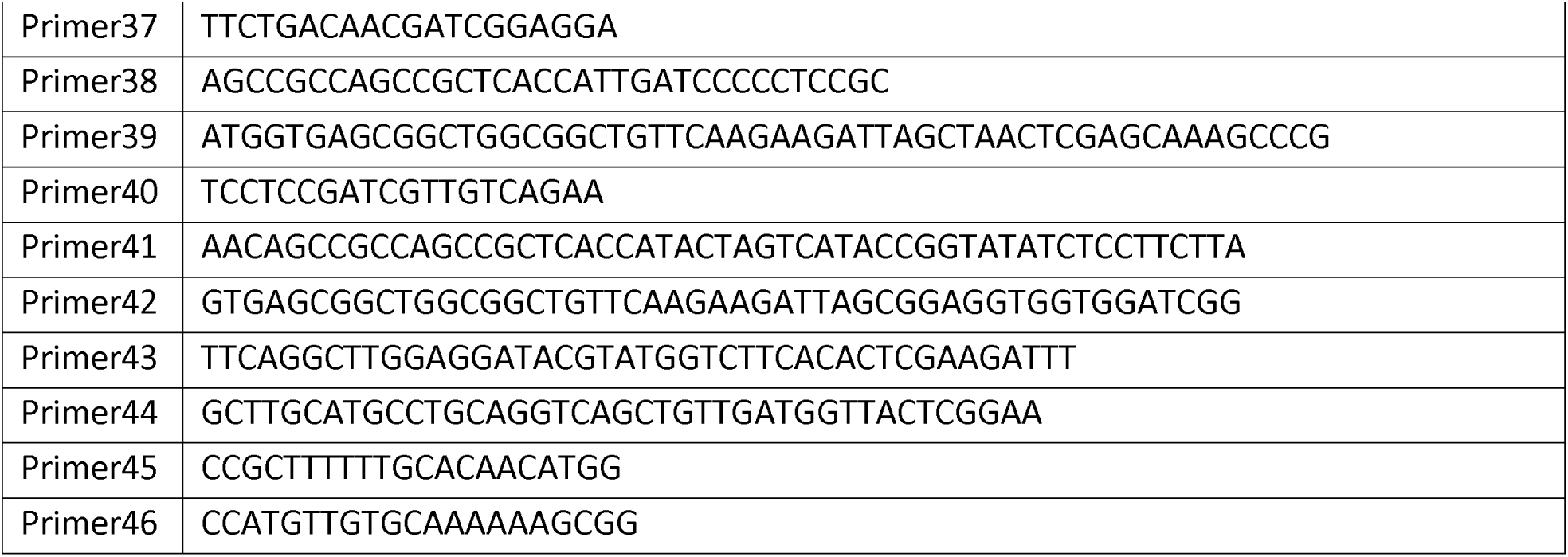
Primer sequences.

**Table S2.**
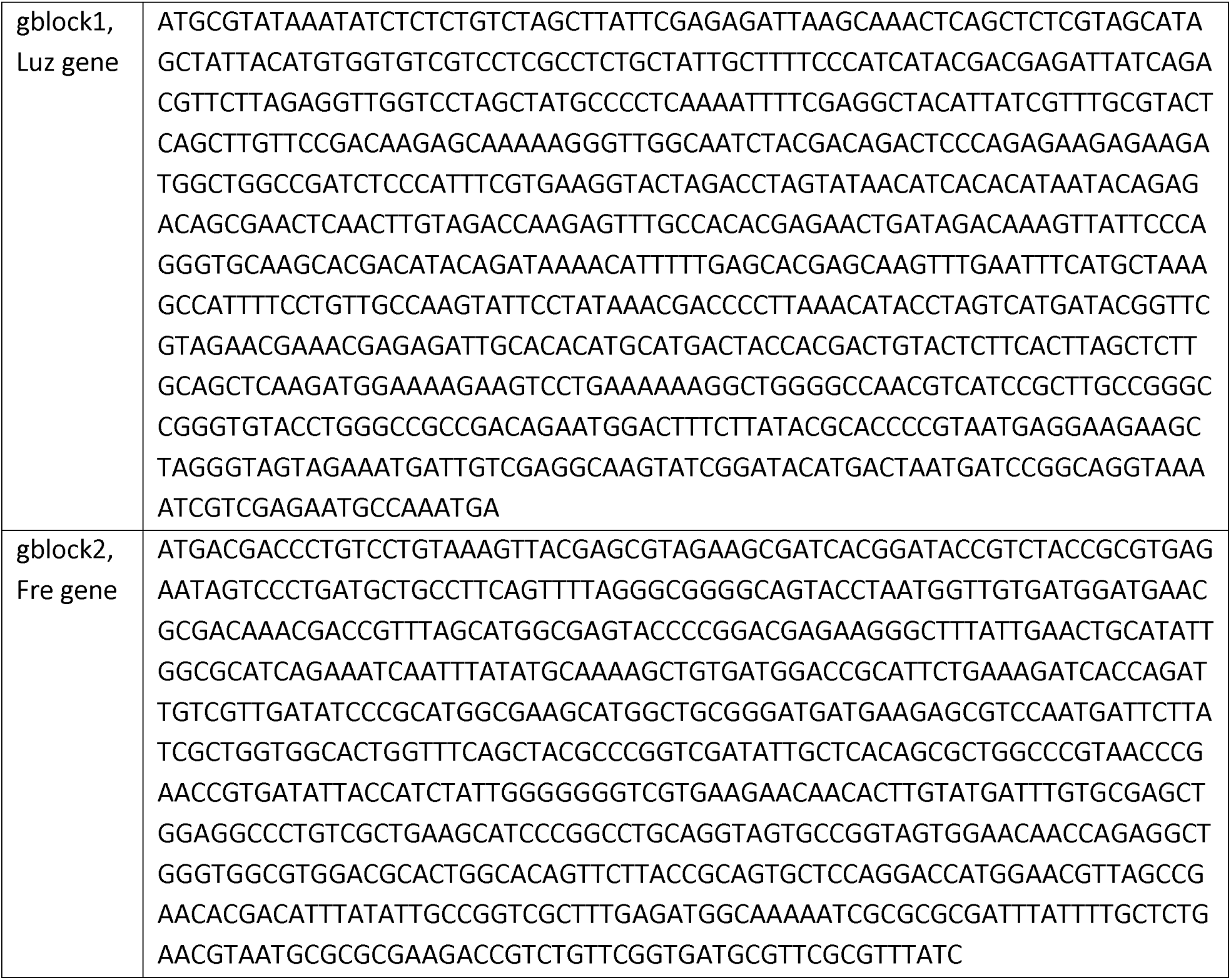
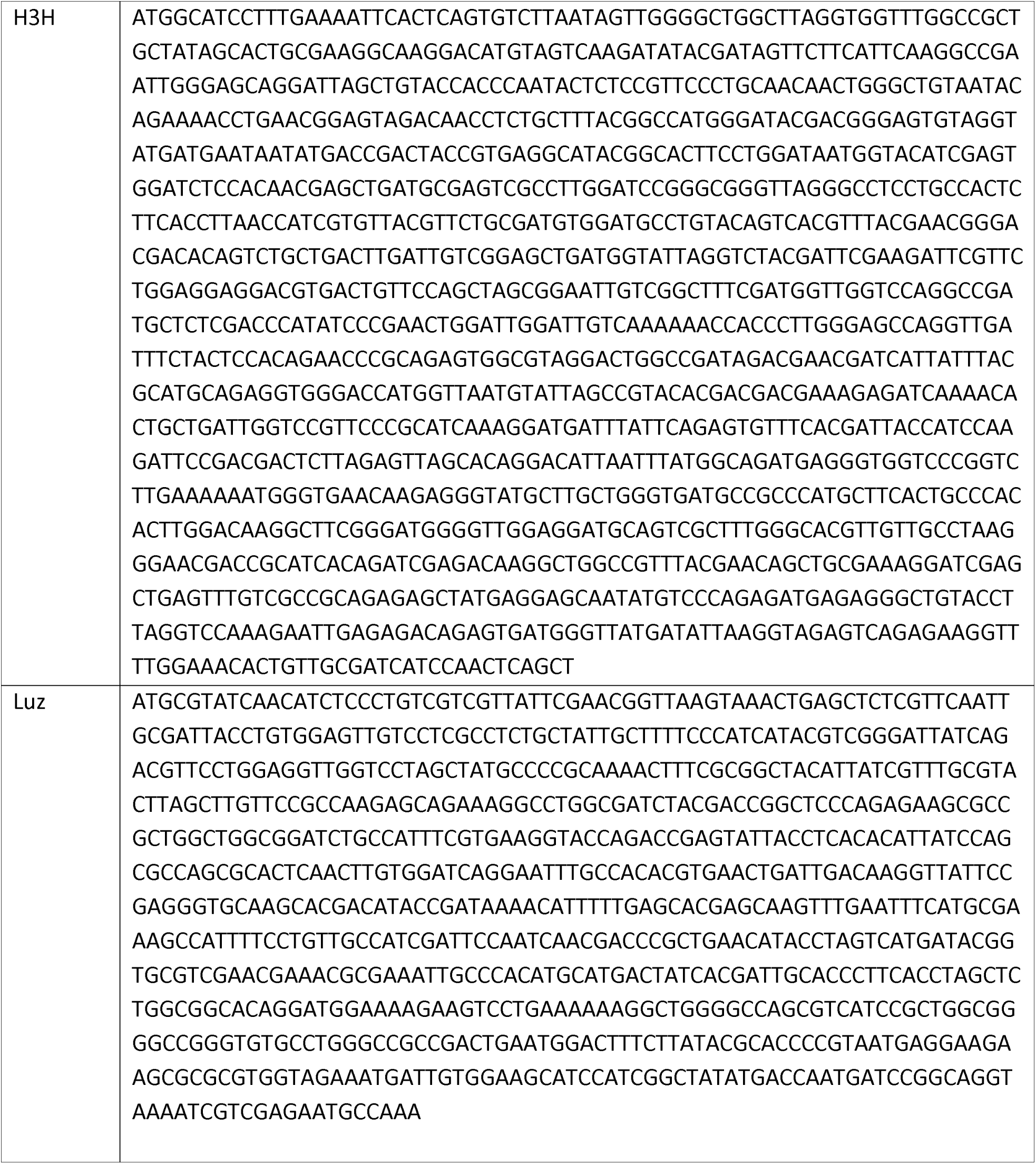
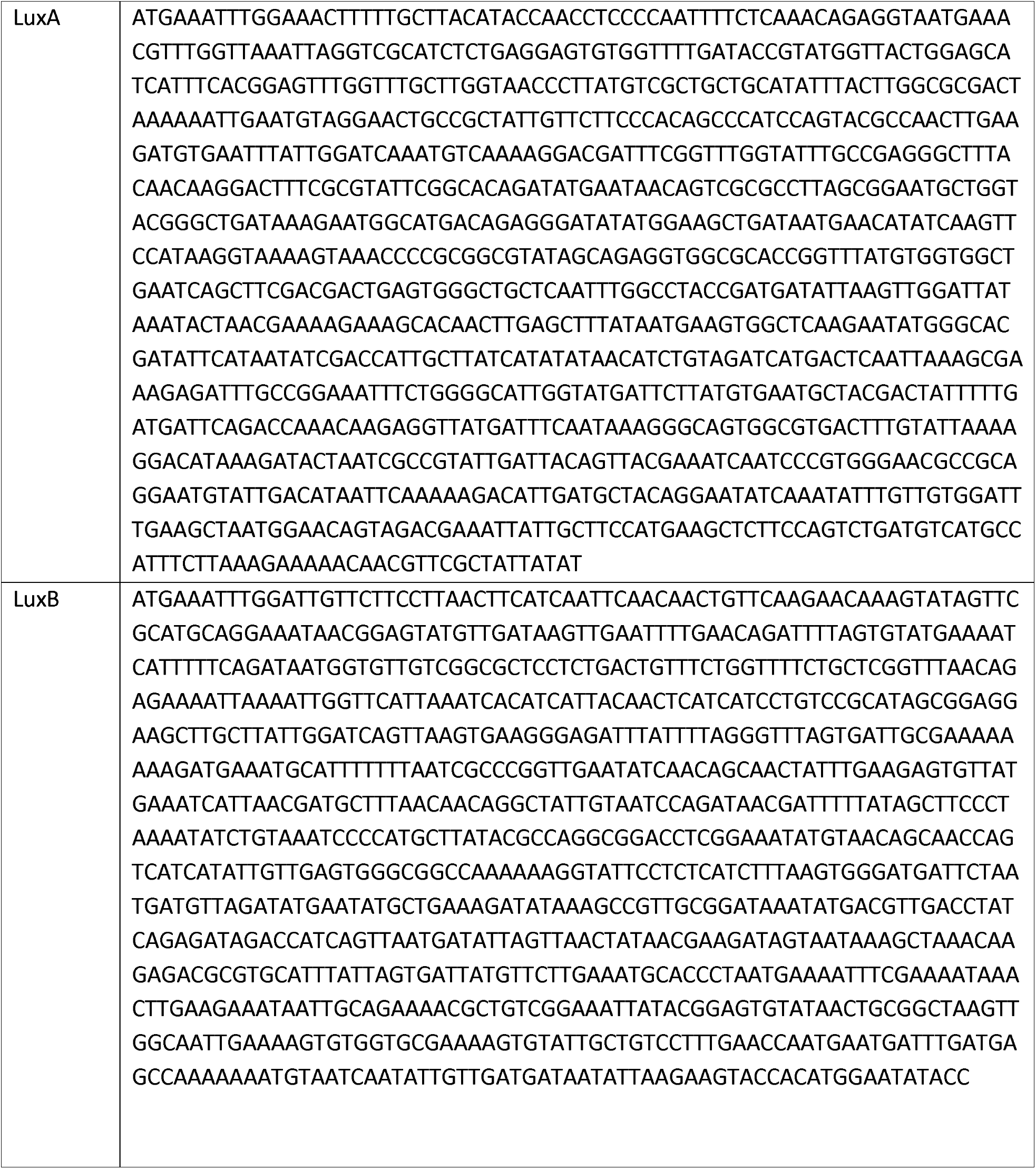
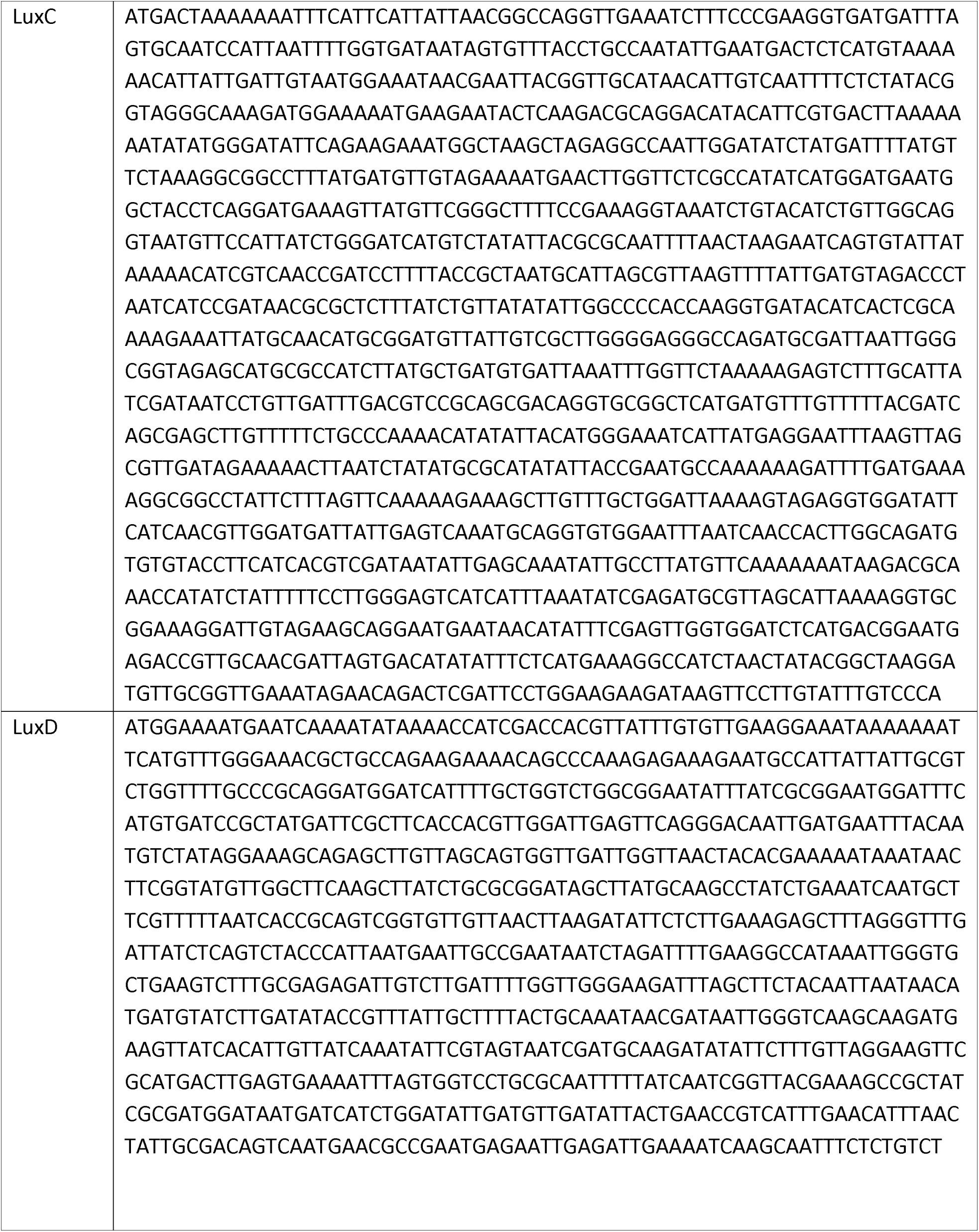
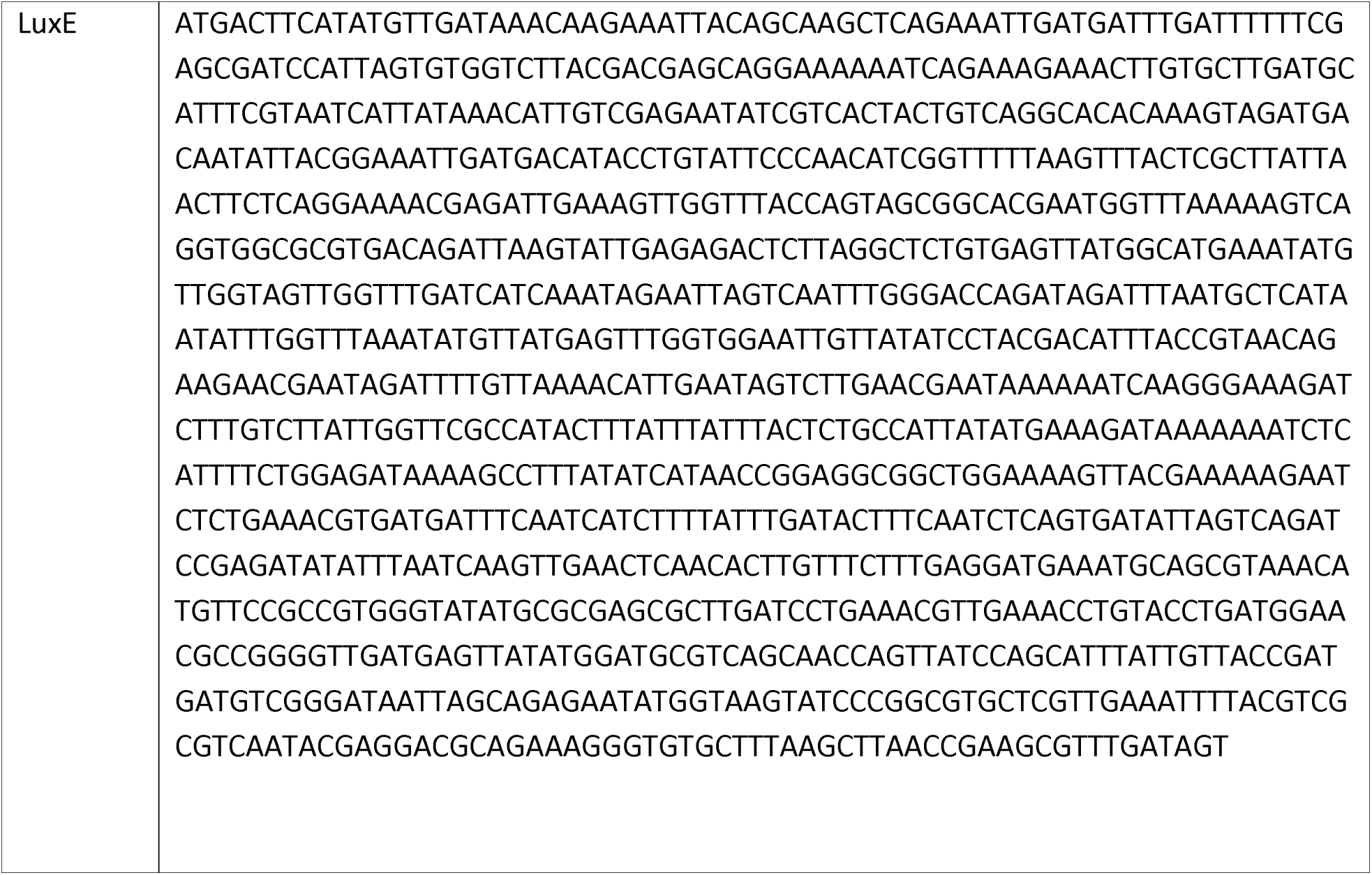
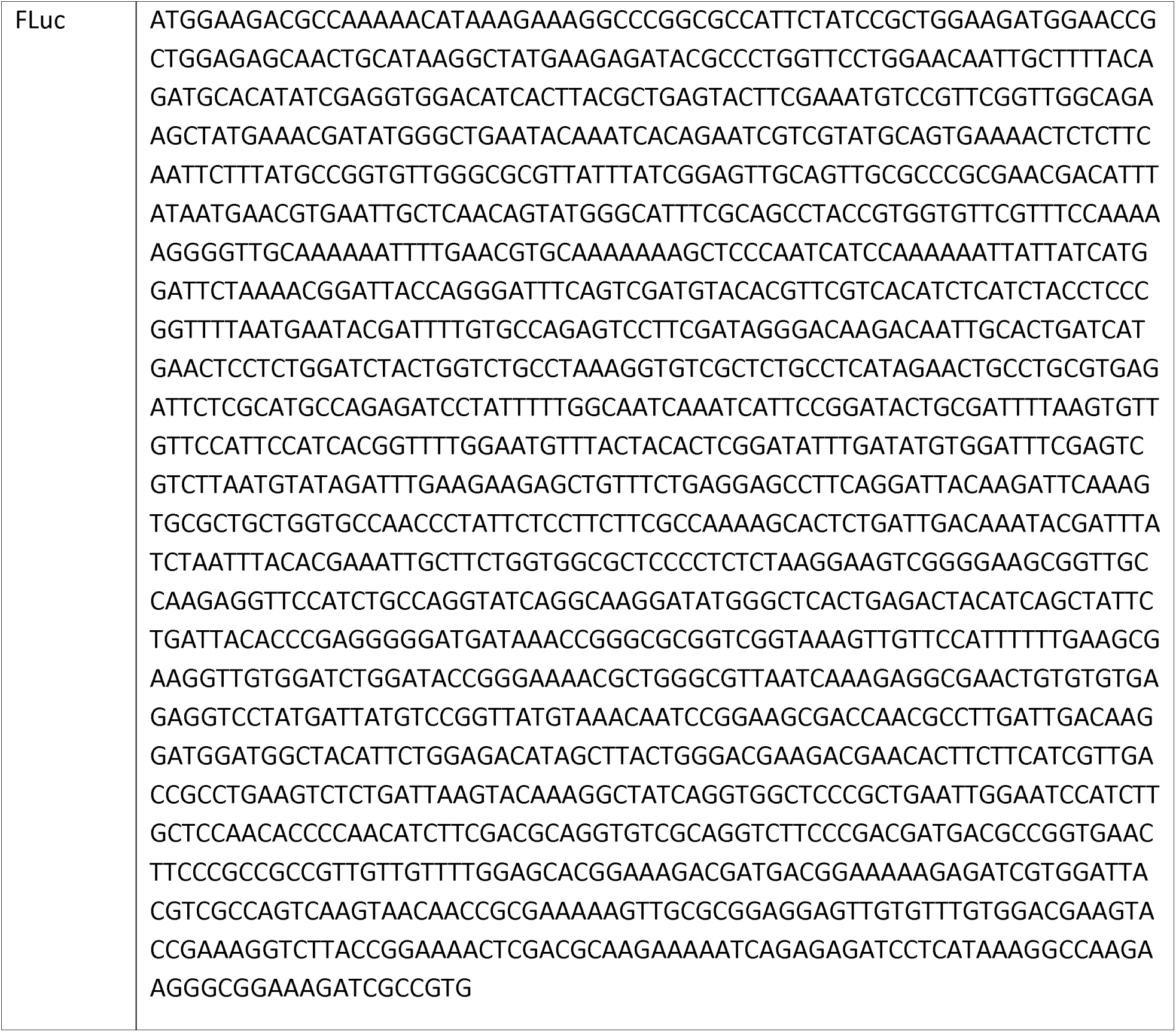
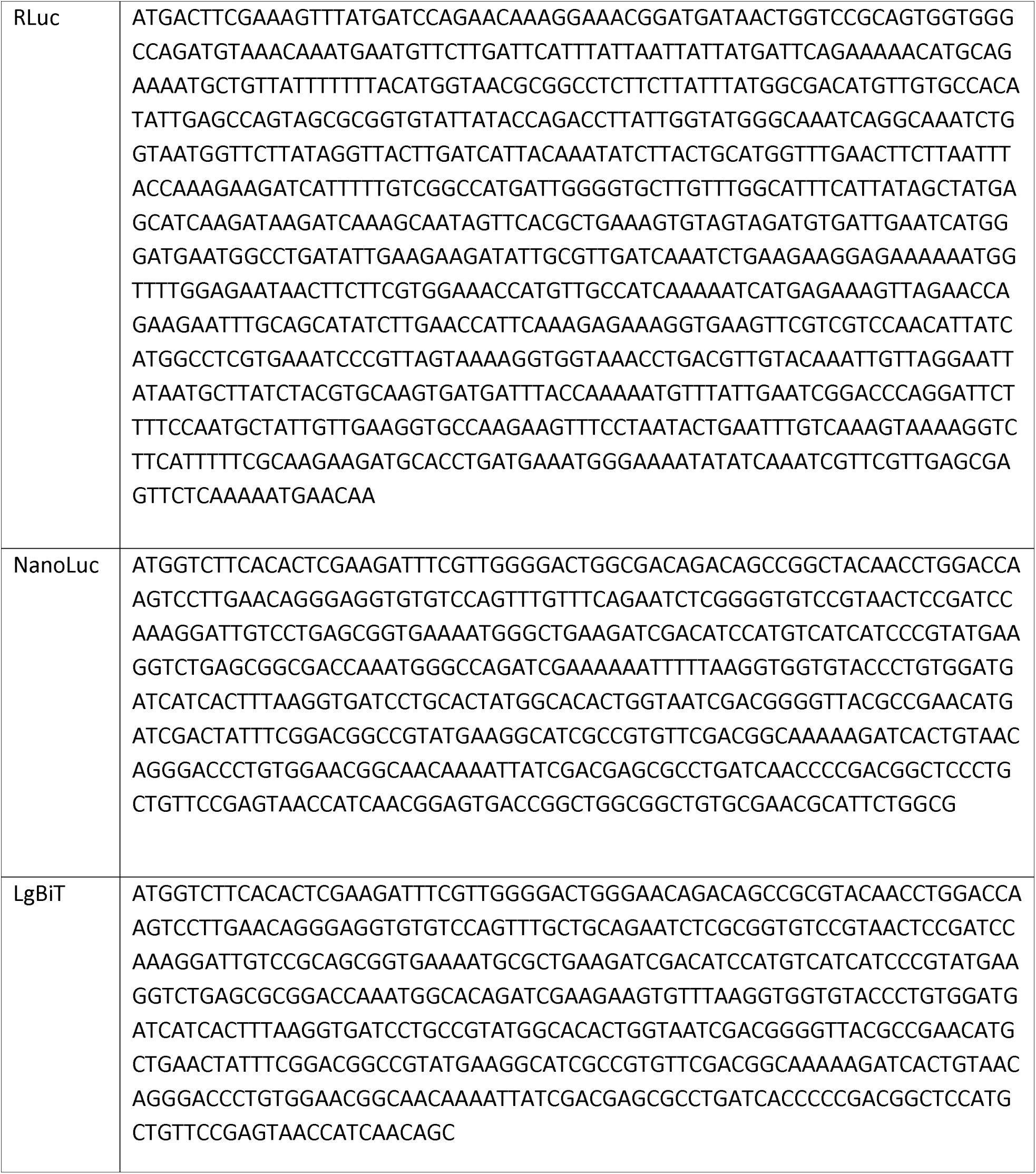
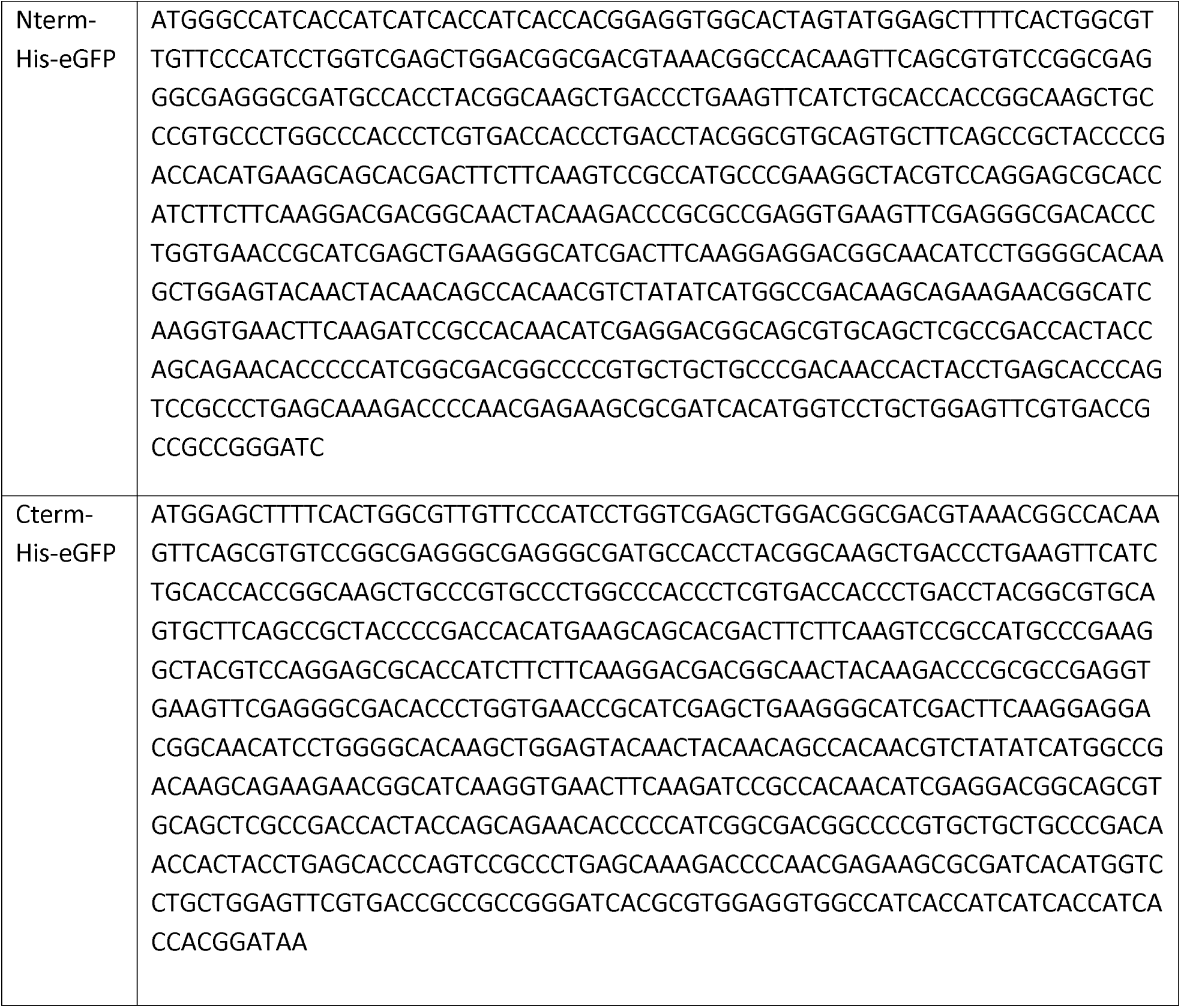
Gene sequence information.

**Table S3.** Cloned plasmid information.

| Name | Purpose |
| --- | --- |
| pTXTL-T7max-H3H | H3H TXTL expression |
| pTXTL-T7max-Luz | Luz TXTL expression |
| pTXTL-T7max-LuxA | LuxA TXTL expression |
| pTXTL-T7max-LuxB | LuxB TXTL expression |
| pTXTL-T7max-LuxC | LuxC TXTL expression |
| pTXTL-T7max-LuxD | LuxD TXTL expression |
| pTXTL-T7max-LuxE | LuxE TXTL expression |
| pTXTL-T7max-Fre | Fre TXTL expression |
| pTXTL-T7max-FLuc | FLuc TXTL expression |
| pTXTL-T7max-RLuc | RLuc TXTL expression |
| pTXTL-T7max-NanoLuc | NanoLuc TXTL expression |
| pTXTL-T7max-HisGFPLuz | N-terminal His-tag eGFP fused Luz TXTL expression |
| pTXTL-T7max-LuzGFPHis | C-terminal His-tag eGFP fused Luz TXTL expression |
| pTXTL-T7max-HisGFPHiBiT | N-terminal His-tag eGFP fused HiBiT TXTL expression |
| pTXTL-T7max-HiBiTGFPHis | C-terminal His-tag eGFP fused HiBiT TXTL expression |
| pLumi-H3H | H3H carrying Rosetta 2 extract preparation |
| pLumi-LgBiT | LgBiT carrying Rosetta 2 extract preparation |

